# A framework for evaluating edited cell libraries created by massively parallel genome engineering

**DOI:** 10.1101/2021.09.23.458228

**Authors:** Simon Cawley, Eric Abbate, Christopher G. Abraham, Steven Alvarez, Mathew Barber, Scott Bolte, Jocelyne Bruand, Deanna M. Church, Clint Davis, Matthew Estes, Stephen Federowicz, Richard Fox, Miles W. Gander, Andrew D. Garst, Gozde Gencer, Andrea L. Halweg-Edwards, Paul Hardenbol, Thomas Hraha, Surbhi Jain, Charlie Johnson, Kara Juneau, Nandini Krishnamurthy, Shea Lambert, Bryan Leland, Francesca Pearson, J. Christian J. Ray, Chad D. Sanada, Timothy M. Shaver, Tyson R. Shepherd, Joshua Shorenstein, Eileen C. Spindler, Craig A. Struble, Maciej H. Swat, Stephen Tanner, Tian Tian, Ken Wishart, Michael S. Graige

## Abstract

Genome engineering methodologies are transforming biological research and discovery. Approaches based on CRISPR technology have been broadly adopted and there is growing interest in the generation of massively parallel edited cell libraries. Comparing the libraries generated by these varying approaches is challenging and researchers lack a common framework for defining and assessing the characteristics of these libraries. Here we describe a framework for evaluating massively parallel libraries of edited genomes based on established methods for sampling complex populations. We define specific attributes and metrics that are informative for describing a complex cell library and provide examples for estimating these values. We also connect this analysis to generic phenotyping approaches, using either pooled (typically via a selection assay) or isolate (often referred to as screening) phenotyping approaches. We approach this from the context of creating massively parallel, precisely edited libraries with one edit per cell, though the approach holds for other types of modifications, including libraries containing multiple edits per cell (combinatorial editing). This framework is a critical component for evaluating and comparing new technologies as well as understanding how a massively parallel edited cell library will perform in a given phenotyping approach.

## Introduction

Genome engineering methodologies are transforming biological research and discovery. Approaches based on CRISPR technology have been broadly adopted due to the relative ease of targeting defined genomic regions using specific guide RNAs (gRNAs) (Jinek et al. 2012). While there has been a large focus on modifying one or a small number of sites for translational research and therapeutics, there is growing interest in the generation of massively parallel edited cell libraries (Ding et al. 2014; Frangoul et al. 2020; Wilkinson et al. 2021). These libraries can accelerate the pace of genome discovery or cell engineering by allowing for the simultaneous interrogation of hundreds to thousands of loci in a single experiment. Current genome-wide approaches typically either leverage knock-out libraries – largely relying on error-prone repair processes for sequence disruptions – or rely on transcriptional modulation by tethering a nuclease-deficient Cas9 with a transcriptional repressor or activator to modulate gene expression (Mali et al. 2013; Cong et al. 2013; Gilbert et al. 2014). Recently, the generation of genome-wide libraries of precise edits has been described in microbes and human (Garst et al. 2017; Sadhu et al. 2018; Bao et al. 2018; Sharon et al. 2018; Hanna et al. 2021). This ability to make more refined changes will provide greater precision and information around genotype-phenotype relationships. Comparing the libraries generated by these varying approaches is challenging and groups typically take different approaches and measures in reporting their work. What is currently lacking is a common framework for defining and assessing the characteristics of these libraries.

The evaluation of these complex libraries can be challenging. The library represents a mixed population, with some cells containing the desired edit and the remaining cells constituting a Burden Population (Table 1) of cells containing incomplete, unintended or no edits. The population of cells containing the designed edits will also be a mosaic, with individual edit representations being driven by the representation of the design in the reagent pool, the functionality of the guide, the edit rate at different loci and any fitness effects an edit may have on an individual cell. Frequently the efficiency of massively parallel editing experiments is extrapolated based on experiments where editing has been performed in isolates rather than in a pooled manner (Sadhu et al. 2018; Sharon et al. 2018). Although this methodology is more experimentally tractable, it is not necessarily predictive of performance in a pooled setting. Additional biological factors can strongly affect outcomes, such as differential growth rates of cells that have undergone the editing process, the introduction of edits that impair cell viability to varying degrees, cells in which no double-stranded break (DSB) is created and which thus grow faster, and cells in which a DSB is created with failure to repair leading to their depletion. All of these factors impact the final library composition. In general, it is preferable for a library to contain a high fraction of edited cells, with an even representation of edits. Understanding the library composition is critical for assessing if a cell library is fit for a given phenotyping regime, though in practice obtaining this information can be technically challenging or cost prohibitive.

**Table 1:** terms and definitions useful for characterizing complex cell libraries

| TERM | DEFINITION |
| --- | --- |
| **BURDEN POPULATION** | The population of cells in a library that is either unedited or contains unintended edits. |
| **COMPLETE INTENDED EDIT** | A precise edit that includes all modifications specified in the repair template (sometimes referred to as the homology arm) with no additional unintended modifications (Figure 1). |
| **EDIT COEFFICIENT OF VARIATION (EDIT CV)** | An aggregate measure across all the edits in a library, the coefficient of variation for the frequencies of the Complete Intended Edits in the edited cells of the library, defined as the standard deviation of edit frequencies normalized to their mean. |
| **EDIT FRACTION** | The fraction of cells in a library containing the Complete Intended Edit at the locus of interest (in a precise editing library) or an edit in the target region (in an imprecise editing library). |
| **EDIT FRACTIONAL RICHNESS** | The Edit Richness (see below) scaled by the library size, a value in the range [0, 1]. |
| **EDIT RICHNESS** | The number of unique Complete Intended Edits present in a sample. |
| **INTENDED EDIT** | The modification of specific bases in a defined region of a genome.<br>(https://www.nist.gov/programs-projects/nist-genome-editing-lexicon#3.4) |
| **REAGENT COEFFICIENT OF VARIATION (REAGENT CV)** | An aggregate measure across all the editing reagents in a library, the coefficient of variation for the frequencies of the editing reagents (typically plasmids, episomes or virus) in the library. Defined as for Edit CV |
| **REAGENT FRACTIONAL RICHNESS** | The Reagent Richness (see below) scaled by the library size, a value in the range [0, 1]. |
| **REAGENT RICHNESS** | The number of unique reagents present in a sample. |
| **SCREENER'S SCORE** | The predicted Edit Fractional Richness for a 1x screen (number of isolates screened = number of designs in library) assuming a 30% Edit Fraction. |
| **SELECTOR'S SCORE** | The predicted Edit Fractional Richness for a selection assuming $1 \times 10^6$ cells and 30% Edit Fraction. |
| **TRACKABILITY** | The ability to identify the reagents associated with a cell in a complex cell library. |

Here we describe a framework for evaluating massively parallel libraries of edited genomes based on established methods for sampling complex populations. We define specific attributes and metrics that are informative for describing a complex cell library and provide examples for estimating these values. Obtaining all of these measures may be challenging or expensive, so *we* also provide a theoretical framework to allow assessment of a given library in the absence of some desired data points. We also connect this analysis to generic phenotyping approaches, using either pooled (typically via a selection assay) or isolate (often referred to as screening) phenotyping approaches. We approach this from the context of creating massively parallel, precisely edited libraries with one edit per cell, though the approach holds for other types of modifications, including libraries containing multiple edits per cell (combinatorial editing). This framework is a critical component for evaluating and comparing new technologies as well as understanding how a massively parallel edited cell library will perform in a given phenotyping approach.

## Library Characterization

Massively parallel genome engineering results in a library of cells, where most cells contain design reagents (that is, the combination of gRNA and repair template) encoding distinct edits. Each design reagent is represented in hundreds to thousands of cells. In microbial libraries, these reagents are often maintained as plasmids, while in mammalian libraries, episomes or genome-integrating vectors, such as lentivirus, must be used if the reagents are to be maintained within the population over the course of an experiment. In many cases, the reagents are attached to a barcode, or are used as a barcode themselves to track which cells contain specific reagents. If selection pressure is applied to the library, these reagents may also serve as a proxy for genotyping the specific edit. A percentage of the population will contain the desired edits, while the remaining population constitutes a Burden Population. In order to characterize such a library, we must define and measure several characteristics. Table 1 provides a list of terms and measures useful for characterizing libraries.

## Definitions Useful for Library Characterization

### Defining an edit

When using CRISPR-Cas based systems to generate a desired sequence variant through precise editing, a guide and repair template are defined (commonly through software). In many cases, auxiliary edits to the PAM site are included to prevent the nuclease from recutting the edited locus. We define a ‘Complete Intended Edit’ as an instance where the repair template sequence (the desired variant and any auxiliary edits) is faithfully and completely placed into the genome (Figure 1) and no other changes occur in the genome. Cases where only part of the repair template sequence is conferred to the genome are classified as incomplete edits and are considered part of the burden, though there will be differences from the reference sequence. Unintended events (off-target editing, reagent integration, or other large-scale genome rearrangement), either occurring at the edit locus or elsewhere in the genome, are also considered part of the Burden Population along with unedited cells.

**Figure 1.**
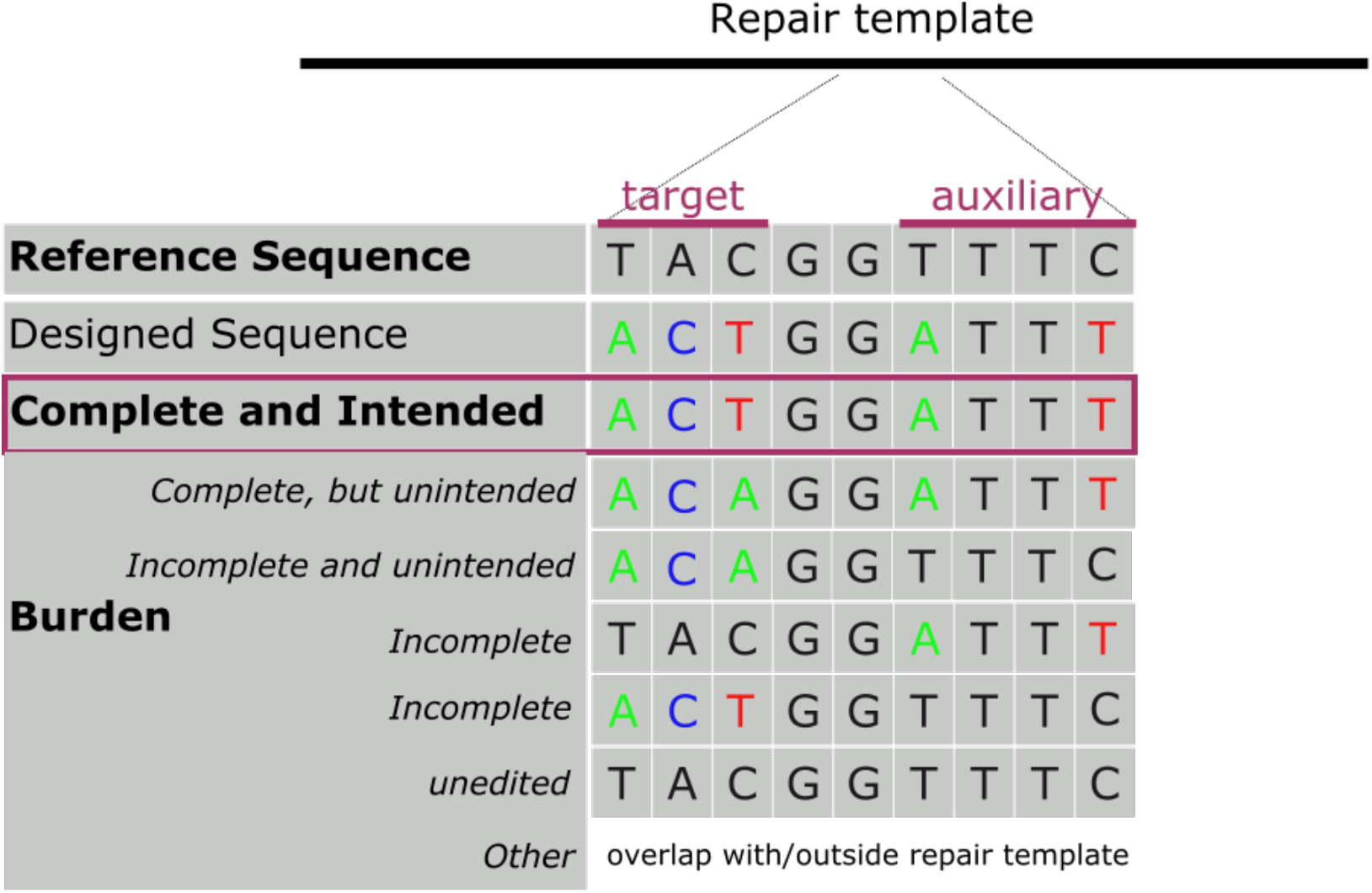
Challenges of edit identification in a large pool of precisely edited cells. A complete and intended edit occurs only when the complete repair template is faithfully placed in the genome; this includes the desired edit and any auxiliary edits made to prevent recutting of the edited locus. Cases where only part of the repair template are incorporated into the genome are considered incomplete and count as burden rather than an edit, even if they include the desired variant. Any other unintended or unedited cells are also considered part of the burden.

When producing imprecise edits, such as in the case of non-homologous end joining (NHEJ)-mediated knockout libraries, the concept of a Complete Intended Edit is not relevant. However, in this case, the desired events would be insertion-deletion events occurring at the target site. Events that do not lead to a true loss of functional protein (knockout) or that happen outside of target region would fall into the Burden Population. In this framework, only Complete Intended Edits (in precise editing) or target site changes leading to a knockout (in imprecise editing) are considered edits. A formal definition of what is meant by an edit allows us to develop a more rigorous framework by which to evaluate these complex cell libraries. In the discussion that follows, the term “edit” refers to Complete Intended Edit unless indicated otherwise.

### Estimation of the Edit Fraction

The Edit Fraction is a critical component of characterizing a massively parallel genome engineered library. Ideally, we would like to identify all edits that occurred within a population. In practice, this is challenging because of the mosaic nature of the library; at any given locus, the count of reference sequence representation will far exceed the count of edit-containing sequences. Fortunately, determination of the overall Edit Fraction does not require complete evaluation of all members of the library. We describe two approaches for identifying the Edit Fraction in a library: a shallow sampling of the library by deeply sequencing isolates or a deeper sampling of the library by shallow sequencing of a pool of cells (Figure 2).

**Figure 2.**
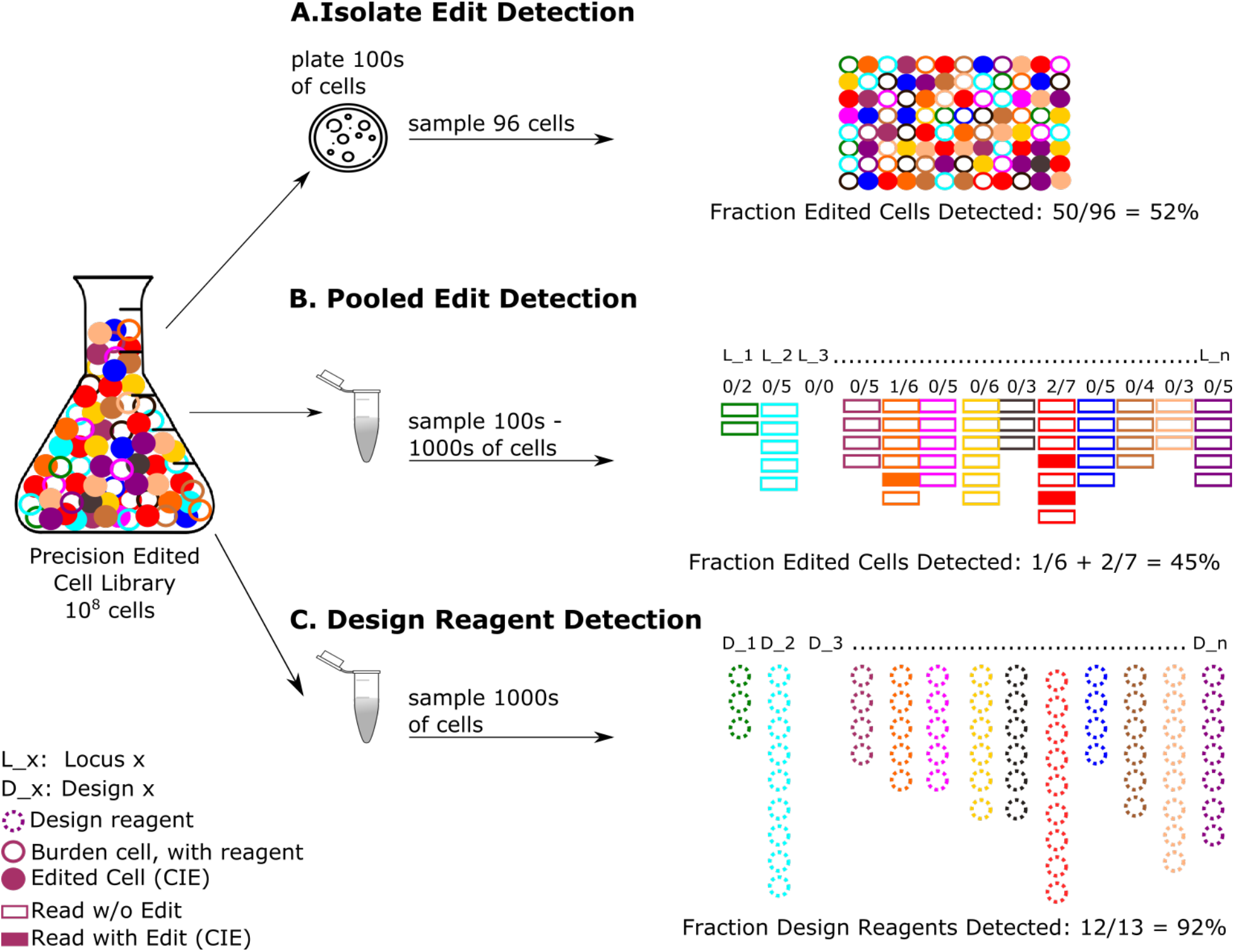
Measurements of interest when evaluating a multiplex precisely edited library. This simplified example is based on a contrived library targeting 13 distinct edits, with half of the cells in the pool containing a Complete Intended Edit and 12 of the designs represented. Open circles represent cells of the Burden Population, most of which will contain editing reagents if selection pressure is maintained or if the trackability reagent is intentionally integrated into the genome. Dashed circles represent the design reagent. Rectangular boxes represent sequence reads, open are wild type while filled are Complete Intended Edit-containing reads. **A.** A shallow library sampling but deep sequencing approach involves edit detection by selecting isolates and performing whole genome shotgun (WGS) analysis. For the isolates selected, this can provide detailed edit data, as well as information on any unintended events, but the approach samples only a small number of cells in the library. It is important to use sufficiently deep sequencing on each isolate to provide good power for detecting edits. **B.** An alternative approach involves doing a broad library sampling but shallow sequence assessment of the library to obtain an estimate of the fraction of cells containing an edit. As with the previous approach, many individual edits that are present in the pool will be absent from the sample; nevertheless, an estimate of Edit Fraction *f* can be obtained by summing the fraction of edited reads at each locus (designated by L_n). At approximately 1000x coverage and with Edit Fraction *f*, 1000*f* edited cells will be sampled. Increasing read depth will increase the number of cells sampled, but very high coverage would be required to deeply assay at each edit locus. **C.** Design distribution can be measured directly from the reagents, typically through a short-read sequencing (NGS) assay using amplification handles. The reagents will be detected in both the edited and Burden Populations, and this assay will not distinguish those populations in the absence of strong selection for edited cells.

One way to assess the Edit Fraction is to sample isolates selected from the population (Figure 2A). After sufficient cell divisions, standard sequencing approaches, such as whole genome shotgun (WGS) of each isolate, can be employed. This requires only collection and growth of isolates (typically by low density plating and picking single colonies into a 96-well plate) and library preparation. While this produces a large number of reads outside of the targeted locus that do not contribute to edit detection, these reads can be assessed for off-target events. Alternatively, one could take an approach to identify the design reagent in each isolate (see below), and then use a targeted sequencing approach, such as hybrid capture or genomic amplification, to confirm the validity of the edit. This approach has the benefit of more efficiently utilizing sequencing reads but takes longer and requires two library preparations, in addition to the creation of custom reagents for each edit locus. Regardless of whether whole genome or targeted sequencing is performed, this isolate evaluation approach generally results in very shallow sampling of a library.

An alternative approach to characterizing the Edit Fraction in a library employs limited WGS on the entire population of cells at a shallow read depth, an approach we term pooled WGS (pWGS) (Figure 2B). While the population of cells used as input for this analysis may number in the millions, the cost of sequencing will typically limit the number of cells ultimately sampled, often in the range of a few hundred to a few thousand. For example, if an experiment involves sequencing to an average genomic coverage depth of 1000x, it will profile approximately 1000 cells’ worth of DNA at each targeted edit locus. In contrast to isolate sampling, the pooled approach limits the manual work of colony isolation and growth at the expense of greater complexity in sequence analysis. If a pWGS assay is tuned to sequence roughly 1000 genomes’ worth of DNA per locus, then for an edit library of 1000 or more members, the assay should be viewed as a sampling of mainly the right tail of the edit frequency distribution. Sampling deeper would require substantially more sequencing, on the order of billions of read pairs or more (Figure 3D and supplemental section 8). Even though the pWGS sampling depth is typically shallow and thus incapable of providing reliable data on a per-design basis, the sum of the per-design Edit Fractions produces a reliable estimate of the overall Edit Fraction in the library (Figure 3A). In either the isolate or pWGS approach, many edits that are present in the pool will be missed in the sequencing results due to being present at very low frequency relative to the per-locus sampling depth. Despite the absence of many of the edits in the sample, making the assumption that the underlying edit frequencies follow a parametric distribution can allow for reliable estimation of the Edit CV (Table 1 and Figure 3D). In situations where the edits are clustered in a subset of the genome, targeted sequencing approaches can provide a more cost-efficient readout of the edit frequencies. Assay replicates will provide differing parameter estimates due to sampling biases in the context of shallow coverage; therefore, inspection of confidence intervals is helpful to guide appropriate interpretation.

**Figure 3:**
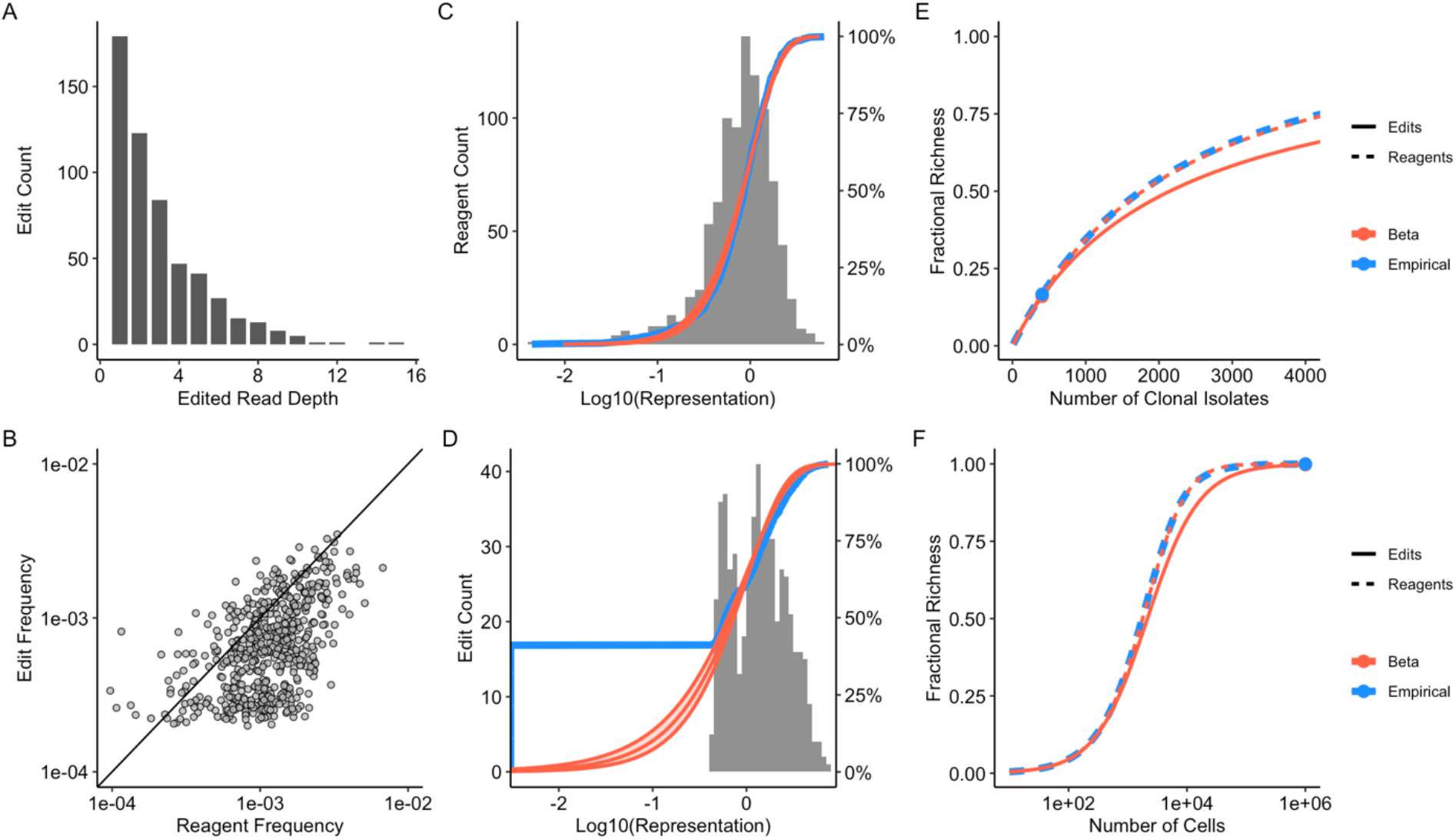
Example usage of pWGS and design reagent amplicon sequencing assays to characterize an E. coli edit library. After exclusion of controls, the library consists of 928 designs including insertions, deletions and substitutions spanning the genome. The resulting edits are not expected to result in any notable effects on cellular fitness. **A**: Number of sequencing reads with exact match to expected edits in a pWGS run. The pWGS run included 157M 2×150 read pairs. After exclusion of reads failing quality filters the mean coverage depth fully spanning the targeted edits is 3434. Summing the per-locus Edit Fractions produces an estimate of 0.44 for the overall Edit Fraction in the pool, thus the pWGS run profiles approximately 1501 genomes’ worth of DNA overall. A total of 1615 edited reads is seen, comprising 546 unique edits (y-axis) with read depth per edit ranging from 1 to 15 (x-axis). **B**: Scatterplot comparing the edit frequencies estimated from pWGS with design reagent frequencies estimated from amplicon sequencing of reagents. **C**: Histogram and cumulative distribution function (CDF) of reagent representation (defined as the product of reagent frequency and library size), measured by amplicon sequencing of the design reagents. The assay consists of 3.0M reads. Fitting the design reagent frequencies to a beta distribution via maximum likelihood estimation (MLE), the data are well described by a beta distribution with mean 1/928 and CV 0.73. **D**: Histogram and CDF as in C, but for the representation of edits as measured by pWGS. Given that the pWGS run is sampling roughly 1501 genomes’ worth of DNA per locus, it should be viewed as a sampling of mainly the right tail of the edit frequency distribution. The fraction of the edit library that is observed at least once is 0.59. Fitting edit frequencies with a beta distribution via MLE, the estimate of CV is 1.01. Observation of a greater fraction of all possible edits in the library would require substantially more sequencing. For example, if the goal were to directly observe 90% of the edits in pWGS, it would require detection of edits whose frequencies among the 44% of edited cells is around the 10th percentile of the reagent frequency distribution, or 1e-4. Aiming for an expected edit read count of 10, to have a reasonable chance of observing edits at the 10th percentile, it would take a mean coverage depth of 213K. This is 62-fold larger than the actual coverage depth for the pWGS run, which would require a total sequencing throughput of 9.8B read pairs. **E**: Screener’s curve, showing the predicted Reagent Fractional Richness (solid curve) and Edit Fractional Richness (dashed curve) as a function of the number of clonal isolates phenotyped in a screening experiment. The red curves are based on a beta binomial model fit. The blue curve is a prediction based on the nonparametric estimate of the distribution of reagent frequencies, a nonparametric fit to the edit frequencies is not useful given the limited sampling depth of the pWGS data. The point indicated on the curve corresponds to the Screener’s score, which is the predicted Edit Fractional Richness when sampling depth is equal to the library size times the Edit Fraction. **F**: Selector’s curve, showing the same data as in E but with the x-axis changed to log scale and domain extended to cover the deep sampling that is typically relevant for the large number of cells sampled in selection applications. The solid point indicated on the curve corresponds to the Selector’s score, which is the predicted Edit Fractional Richness when sampling 1M cells.

### Estimation of Reagent Distribution

Direct detection of edits in massively parallel editing libraries is ideal for assessing library diversity, but in practice it is often prohibitively expensive due to the depth of sequencing required. In lieu of extensive genomic sequencing, many approaches make it relatively straightforward to detect the reagents conferring edits, so profiling the reagent distribution can be a useful proxy for the edit distribution. For microbes, each cell typically contains multiple clonal reagent copies, and most reagents will be present in hundreds to thousands of cells. For mammalian cells, the copy number of the trackable reagent is typically lower, on the order of one to less than 10. Ideally, all designs would be equally represented, but in practice most libraries have a distribution of representation. Every manipulation of the library (reagent manufacturing, transformation, growth of the cell population) introduces an opportunity to alter this distribution. Understanding the distribution of reagents is critical for interpreting phenotyping results and will help define the effect size and significance of results. For example, if a phenotyping approach is assessing depletion of reagents as a measure proxy for genotype (a common approach in essential gene screens), designs in the extreme left tail of the distribution will likely be underpowered for association with a phenotype.

Sequencing the reagent library throughout the experimental process provides useful insight into how various manipulations can impact design reagent distribution. This approach can be useful for approximating edits post-phenotyping, particularly in the case of strong selective pressure. In a library containing a mixture of active and inactive gRNA-donor cassettes, the number of viable edited cells is tightly coupled to gRNA activity, rate of homology directed repair (HDR) and the relative survival rate of edited members of the population. DNA synthesis errors that result in unintended editing events during the homology-directed repair process or poor transformation efficiency can impact uniform representation of intended edits (Roy et al. 2018). These effects can reduce the effective diversity in an edited library, directly impacting the success of phenotyping. For instance, edited variant libraries may lack the desired intended diversity due to editing process failures or takeover by a sub-population of a particular Complete Intended Edit, unintended edits or unedited cells. In each of these cases, the cost and effectiveness of phenotypic investigations will be adversely affected.

Typically, short read sequencing (NGS) of the reagent is used to determine the library distribution from a sample of the library (Fig 2C). Approaches that either detect a barcode (Garst et al. 2017; Sadhu et al. 2018) or the reagents themselves (Bao et al. 2018; Sharon et al. 2018) are used. It is assumed that the read counts for a design reagent are proportional to the number of cells containing that design; thus, a read count is equivalent to a design reagent count. The dispersion of the distribution is measured by the Reagent CV (Table 1, Figure 3C).

Larger Reagent CV values indicate greater variance in the relative abundances of the designs, which can lead to under- or overrepresentation of individual designs. Prior to applying selective pressure, a small Reagent CV is preferable for all phenotyping approaches, though libraries with larger Reagent CVs can still be useful for some experiments. It is important to note that while the Reagent CV is a useful and accessible metric, what matters most for many applications is the **Edit CV** (Table 1). If every design reagent has an equal probability of producing an edit, the Reagent CV and Edit CV will be equal to one another. In most real-world situations there are various sources of bias, including those mentioned above, which result in the Edit CV being larger than the Reagent CV, to an extent that will depend on the experimental context (Figure 3D).

We have introduced measures that can be useful for describing aspects of a massively parallel edited cell library. We next introduce approaches for combining these measures to produce metrics that can be utilized for evaluating these libraries.

## Metrics for Library Evaluation

In this section we define several concepts that utilize the above measurements to provide a fuller characterization of a library. Neither Edit Fraction nor reagent distribution alone can fully characterize the utility of a library. When sampling a library with a high Edit Fraction but poor representation of some or many library members, any phenotyping regime will be continually sampling only a small subset of the desired variation. Alternatively, even representation of the designs with a poor Edit Fraction will lead to over-sampling of the Burden Population. Different phenotyping approaches will be more or less tolerant to deviations in either Edit Fraction or design reagent distribution. Below, we describe metrics that combine these two measures into a score that can be used to quickly assess the utility of a given library.

### Edit Library Richness

When sampling cells or isolates from an engineered cell library, the quantity that is typically most important is the number of unique edits represented in the sample. Borrowing from the ecological literature, the term “richness” is used to refer to the number of unique edits in the sample from the library (Levin et al. 2012). The expected richness *μ_m_* of a sample of m cells or isolates from a library of S edits can be predicted given *f*, the fraction of cells that contain an edit, and the frequencies *p_i_* of each edit among the edited cells.

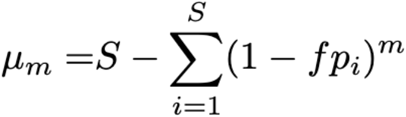

As with other measures, the variance of the sample’s richness can be calculated (supplemental section 1). For some approaches, a variant will need to be observed more than once to provide statistical power for making the genotype-phenotype correlation. In these cases, there is a tractable generalization for when richness is defined in terms of needing at least *n* observations of each edit (supplemental section 2). This is useful in cases where the dynamic range of quantification relies on a set number of observations of the edit. There is an accurate approximation for the mean and variance of richness, useful both for its mathematical convenience and because it reduces computational complexity from *O*(*n*^2^ *S*^2^) to *O*(*nS*) (supplemental section 3).

Under the assumption that all designs have equal probability of conferring their edits, measurements of reagent frequencies and of the Edit Fraction can be used to predict the richness in a variety of circumstances. It is useful to plot the predicted richness against the number of cell isolates evaluated in a screen or selection, producing a “Screener’s Curve” (Figure 3E, 4E and 5C) or a “Selector’s Curve” (Figure 3F, 4F and 5D). These plots serve as a guide to set expectations of what fraction of an edit library will be probed in a screen or selection.

**Figure 4:**
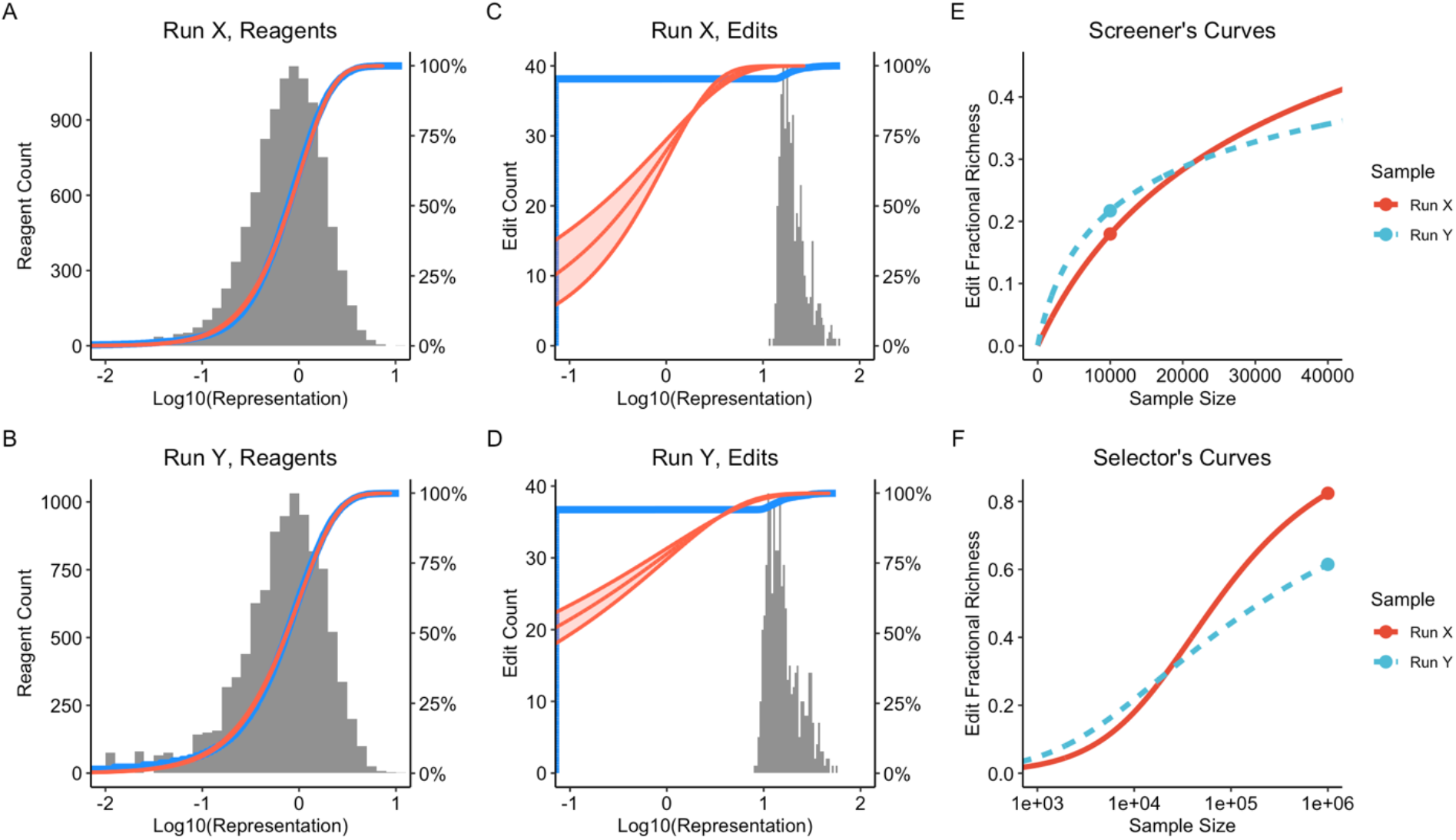
Comparative evaluation of two runs of a 10,000 member *E. coli* library, the runs are named X and Y. **A** and **B**: histogram and CDF (blue) of design frequencies as determined by deep amplicon sequencing of the reagents. The red curves correspond to beta distributions fit by Maximum Likelihood Estimation (MLE). The estimates for Reagent CV are 0.79 and 0.90 for runs X and Y respectively. **C** and **D**: histogram and CDF (blue) of genomic edit frequencies as determined by pWGS. The red curves are beta distributions fit by MLE, the shaded area spans the 95% confidence interval for the edit CV estimates. The estimated edit CVs are 1.54 and 2.48 for runs X and Y respectively. The pWGS assay is a shallow sampling of edits, with an estimated sampling depth of 488 and 724 in runs X and Y respectively, which is very small compared to the library size of 10,000. The pWGS assay also enables estimation of Edit Fraction, the estimates are 0.25 and 0.57 for runs X and Y. Run X has a lower Edit Fraction but also a lower edit CV compared to run Y, so determination of which run is better to use in downstream applications will depend on the situation. **E**: Screener’s curves plotting predicted Edit Fractional Richness against sample size for the two runs. The points on the curves correspond to the Screener’s Scores using the estimated Edit Fractions. For a screen of 20,000 or fewer isolates (twice the library size), run Y is predicted to yield greater Edit Fractional Richness, with its larger Edit Fraction making up for its larger edit CV. **F**: Selector’s curves, like E but with the x-axis expanded to span a range more typical for a selection application. The points on the curves denote the Selector’s Scores, the predicted Edit Fractional Richness when sampling 10^6^ cells. The lower edit CV of run X makes it a better choice for a selection application, despite it having less than half the Edit Fraction of run Y.

**Figure 5:**
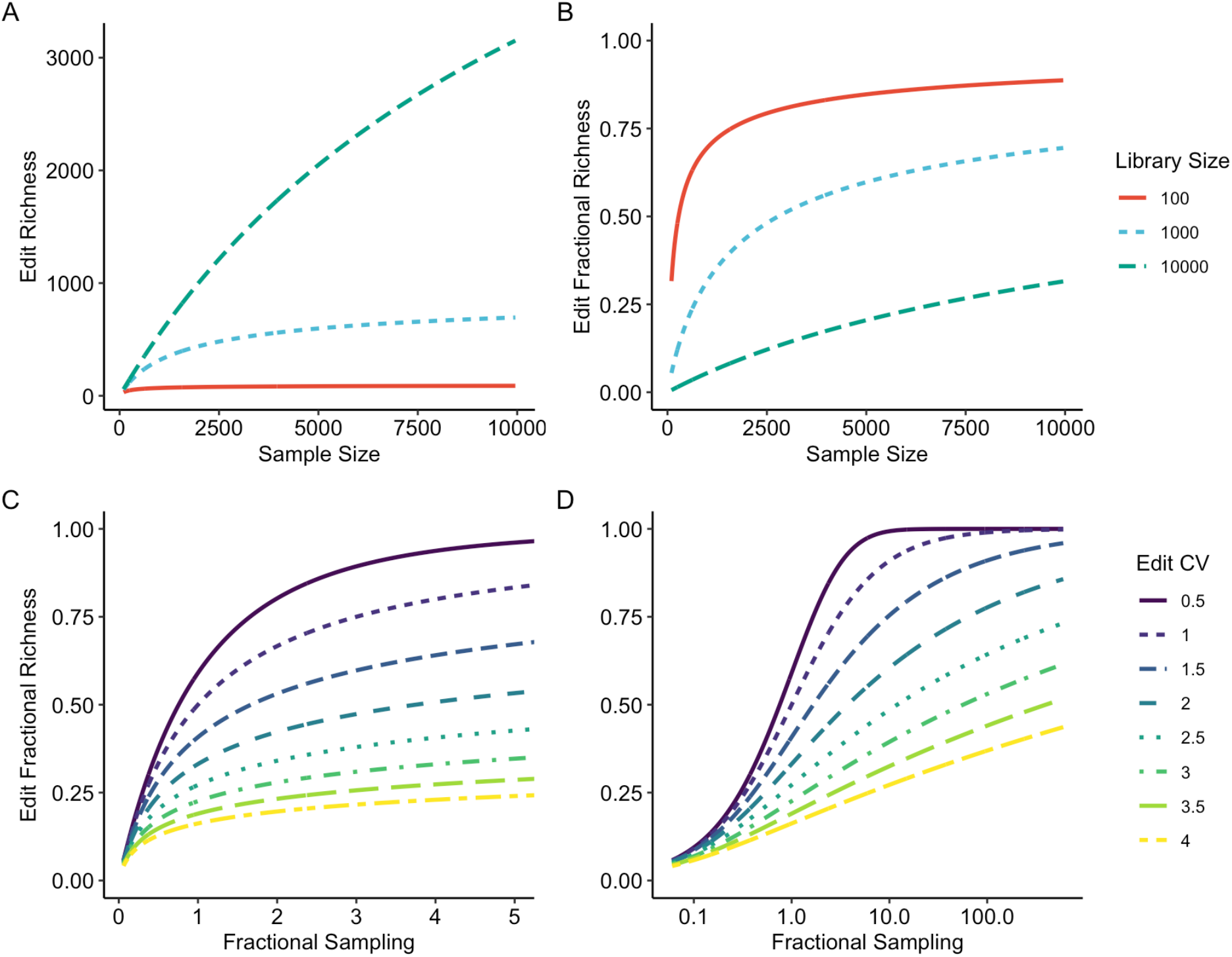
Exploration of richness under the assumption that edit frequencies follow a beta distribution. **A**: Edit Richness for different library sizes, assuming an Edit CV of 1.5 and an Edit Fraction of 0.6. **B**: Edit Fractional richness for the same scenarios as used in A. **C**: Screener’s curves, showing Edit Fractional Richness as a function of Fractional Sampling, with different values for edit CV. Fractional Sampling is defined as the product of sampling depth (the number of cells or isolates sampled) and Edit Fraction divided by the library size. Fractional Sampling and Edit CV are all that is required to predict Edit Fractional Richness under the beta assumption. **D**: Selector’s curves, which are the same figure as C with a log-scale x-axis to enable prediction of Edit Fractional Richness with the deep sampling that is typically used for a selection experiment

The appropriate sample size *m* from which to make richness predictions will depend strongly on the particular situation. In some cases, the cost of phenotyping each sample is high, and the sample size needs to be kept small for practical reasons. In other cases, deep sampling is affordable, and many cells can be sampled. To be able to quantify a library’s suitability for screening and selection applications, and to be able to do so in the absence of an estimate of Edit Fraction, two metrics are introduced - the Screener’s Score and the Selector’s Score. The Screener’s Score is defined as the expected Edit Fractional Richness when sampling *S* times (a 1-fold sampling of the library) and with Edit Fraction set to 0.3. The maximum possible value for the Screener’s Score is 1 – *e*^−0.3^ or 0.26 (supplemental section 4). The Selector’s Score is defined as the expected Edit Fractional Richness when sampling 10^6^ times (a reasonable number of input cells for a selection protocol), with the same Edit Fraction of 0.3. The Selector’s Score can take on any value in the range [0,1]. These scores are intended to be general measures and more detailed information concerning the Edit Fraction would make this estimate more accurate. Figure 4 illustrates how these concepts can be used to quantitatively assess different libraries for screening and selection purposes.

When an estimate of Edit Fraction is available to complement the estimates of design reagent frequencies, the Empirical Screener’s Score and Empirical Selector’s Score can be evaluated in a similar manner, replacing the fixed assumption of 0.3 Edit Fraction with the empirically determined estimate (Figure 3D). These curves aid in understanding the best phenotypic approaches to take given various library characteristics and experimental goals.

### Maximizing Library Richness

The four variables appearing in the expression for richness motivate different approaches for maximizing the richness of a sample, though in practical applications some of the approaches may be inaccessible (supplemental section 4). The first approach is the obvious one of increasing the sample size – the larger the sample, the greater the richness. The second approach is to increase the probability f that a design reagent confers an edit - something that can be achieved, for example, by improving models for gRNA design. The third approach is to increase the library size S. Lastly, the edit CV has a direct impact, with more evenly distributed libraries resulting in greater richness.

For a sample of size m from a library of size S with Edit Fraction f, the maximum richness possible is 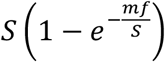, attained for a perfectly even library where all design reagent frequencies are equal to 1/*S* (supplemental section 4).

### Predicting Library Richness

The predictor of library richness introduced above requires an estimate of the frequency of every member of the library. In some situations where deep sampling from the library is feasible it will be possible to get good frequency estimates, but for large libraries it is often desirable to be able to predict richness from shallow sampling, to help guide decisions about when to proceed with deep sampling.

The problem of predicting future richness from an initial sampling is commonly referred to as the unknown species problem in ecology, one of the earliest solutions was the Good-Toulmin estimator (Good and Toulmin 1956). The Good-Toulmin estimator is a nonparametric approach which works well for predicting up to twice the depth as available in the initial sample but beyond that it becomes unstable. An improved nonparametric approach introduced the use of rational function approximations to produce stable estimates at sampling depths orders of magnitude larger than the initial sample (Daley and Smith 2013) and subsequent work extended the approach to predict richness when requiring more than one observation of each library member (https://arxiv.org/pdf/1607.02804.pdf).

An alternative approach is to assume a parametric model to describe the library frequencies. A benefit of the parametric approach is that it can produce good estimates from shallow sampling, as long as the model is a good fit for the underlying data. The beta distribution, described by two parameters, is a natural model to consider and one that is often an excellent fit for genome editing libraries (Figures 3, 4, S4). When using a model for design reagent frequencies where the total library size is known, a constraint is needed to ensure that the frequencies sum to 1, or equivalently, to ensure their mean is 1/S; as a result, there is only one free parameter. It turns out to be convenient to use the CV as the free parameter. When design reagent frequencies follow a beta distribution, there is a closed-form solution available for the expected Edit Fractional Richness, where Edit Fractional Richness is defined as the Edit Richness scaled by the library size (supplemental section 6). For a beta model, Edit Fractional Richness depends on only two parameters - the CV of the design reagent frequencies c, and the sampling fraction *F*, defined as mf/S, which can be thought of as the effective fraction of the library that is profiled in a sampling of *m* cells (Figure 5). The expected Edit Fractional Richness *μ_m,n_* where at least n observations of an edit are required, is well approximated as

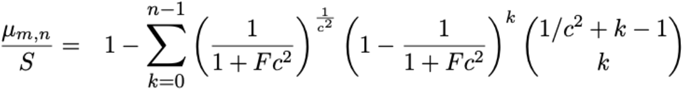

Consistent with the expression for Edit Fractional Richness, the number of observations of each edit in the sample follows a negative binomial distribution with failure probability set to 1/(1 + *Fc*^2^) and failure count set to 1/*c*^2^. There is also an expression for the variance of richness (supplemental section 6). These expressions can be used with the delta method to account for uncertainty in the estimates of CV and Edit Fraction, enabling construction of confidence intervals for Screener’s and Selector’s curves.

Supplemental section 9.3 presents a comparison of parametric and nonparametric estimators of richness on some empirical data.

## Applying These Estimates and Metrics

Massively parallel genome engineered libraries provide rich diversity for a variety of applications. The framework described above can be applied to experimental design, library evaluation and comparing results from different approaches. Below, we describe using this framework to evaluate libraries for utility in either forward engineering or genome discovery applications.

### Forward Engineering Experiments

Forward Engineering of biological systems relies on effective methods to generate beneficial genetic diversity to provide the fuel for evolutionary optimization (Fox and Giver 2011). Screening of isolated genetic variants that drive improved phenotypes becomes an exercise in maximizing richness while managing sampling depth. As noted above, increasing the library size is a way of maximizing richness. Shallow screening of large libraries has proven to be an efficient way to maximize the beneficial diversity rate, as most of the genotypes observed are likely to be unique at lower sampling depth (Alvizo et al. 2014).

The effects of library size, Edit Fraction and Edit CV for screening experiments is shown in Figure 5. The discovery rates for libraries with differing Edit CVs are plotted, showing the effect to which libraries with higher variance in the distribution of the population forces much deeper screening in order to continue to observe unique variants. For forward engineers seeking simply to maximize the discovery rate of beneficial diversity, a shallow sampling from a large library is a particularly effective approach. For shallow sampling, the impact of Edit CV on Edit Fractional Richness is modest, as few of the sampled variants are duplicates. Conversely, with deeper sampling (where researchers desire observing the highest fraction of designs) the effect of a larger Edit CV becomes more limiting. As the Edit CV of the library population increases, it becomes increasingly difficult to observe those designs present at the lower frequencies in the population. Edit Fraction has a linear effect on screening outcomes - halving the edit rate while doubling the sample size results in no net change in expected richness.

## Genome Discovery

While forward engineering is driven largely by the identification of desired phenotypes, genome discovery is often focused on testing specific variants to determine if they drive a phenotype. In this case, a researcher may be more interested in observing all, or most, variants within a library several times in order to develop robust hypotheses around genotype-phenotype correlations. In this case, maximizing library coverage may be the most beneficial approach. When employing an isolate phenotyping approach, this will likely require minimizing library size so that the edits can be sampled multiple times. When employing a selection strategy, increasing library size may be appropriate if Edit CV is held low. This will be driven by the number of times a researcher wants to observe edits in the left tail of the distribution. For more precise genotype-phenotype correlations, assessing more libraries containing a smaller number of edits will likely yield more robust results. Strategic use of the Screener’s and Selector’s Scores in planning experiments can maximize outcomes by informing sampling depth needed to robustly associate genotypic changes with phenotypes of interest.

## Conclusions

As technology continues to improve, the ability to create larger libraries with precise edits will become commonplace. To date, no common standards exist for describing and evaluating cell libraries. This makes comparing libraries produced using different approaches challenging. Perhaps more importantly, a lack of common standards makes planning experiments and evaluating libraries as fit-for-purpose challenging, and these measures differ from lab to lab. Here, we have proposed a framework for evaluating massively parallel libraries of genome engineered cells. We have provided precise definitions around what constitutes an edit. While previous groups have often looked at the reagents within a complex cell library, we demonstrate the value of measuring the fraction of cells within the pool that actually contain an edit and we introduce methodology to directly profile the distribution of edit frequencies. This provides for robust characterization of library properties without needing to employ expensive and labor-intensive approaches to understand editing at every target site. We introduce the concept of edit library richness to more fully describe a library quantitatively, as the Edit Fraction is insufficient to fully characterize a library’s quality. When generating a complex editing library, it is valuable to have a large percentage of the designs represented in the final population, not just have a large Edit Fraction that all contain the same, or a few edits. We also provide models and methods that allow predictions of library quality when some key metrics, typically Edit Fraction, are not available. Development of a robust framework for evaluating complex cell libraries will be necessary to inform which approaches will be useful for phenotypic analysis of a library. Establishment of common methods will facilitate comparing libraries created from various methods. While we have focused on libraries of precise genome edits, the metrics, models and methods proposed here can be applied to any type of library conforming to the general statistical assumptions introduced.

## Supplemental Materials

Mathematical derivations and deeper discussion of the metrics are available in the attached Supplement. Code and data used for analyses can be accessed online at https://github.com/InscriptaLabs/cell_lib_eval_paper

## Acknowledgements

The authors are grateful to Arnold Oliphant, Lior Pachter, and Fritz Roth for their helpful and insightful feedback

## Author Contributions

CGA, SC, CD, ME, SF, RF, MWG, ADG, MSG, ALH, PH, TH, SJ, CJ, KJ, NK, SL, BL, TMS, JS, ECS, CAS, MHS, ST and TT developed the general framework for characterizing a pool of edited cells and created the novel associated metrics. EA, SA, ME, GG, NK, BL, FP, CDS, TRS, and KW used the Onyx^™^ platform to generate the pooled editing data used in this manuscript. MB, DMC, SC, ME, RF, MSG, TH, BL, JCJR, TMS, CAS, and MHS wrote and/or reviewed the manuscript and associated figures. JB, SC, TH, SL, TMS, CAS, MHS, and ST derived the mathematical results in the main text and supplement, and implemented them in bioinformatics pipelines.

