## Supplement for "A framework for evaluating edited cell libraries created by massively parallel genome engineering"

#### 1 Expected Value and Variance of Richness

Consider a collection of $S$ distinct types of objects, in which the frequency of object type $i$ is $p_i$. When sampling from the set, the richness of the sample is defined as the number of unique types of object included in the sample. The fractional richness is defined as the richness divided by $S$. Fractional richness is in the range $[0, 1]$ with a value of 1 corresponding to the inclusion of every possible type of object in the sample.

A term can be introduced to include the concept that each sampled object can be accepted or rejected with a certain probability. Let $f$ be the probability that each sampled object is accepted, conditional on having been included in the sample. As an example of how this can be relevant to genome editing libraries, $p_i$ can represent the frequency of design $i$ in the pool of edited cells and $f$ can represent the probability that design $i$ successfully confers the intended edit.

When sampling $m$ objects without replacement from the set, the richness $R_m$ of the accepted objects in the sample is a random variable whose expected value $\mu_m$ and variance $\sigma_m^2$ can be expressed as

$$\begin{aligned} \mu_m &= S - \sum_{i=1}^{S} (1 - f p_i)^m \\ \sigma_m^2 &= (2S-1)\mu_m - \mu_m^2 - S(S-1) + \sum_{i=1}^{S} \sum_{j \neq i} (1 - f p_i - f p_j)^m \end{aligned} \tag{1}$$

The derivation is outlined in section 10.1.

A more general model would be to allow for object-specific acceptance probabilities $f_i$. In such a scenario, define the overall acceptance probability $f = \sum f_i p_i$ and define $q_i = f_i p_i / f$, then the expected value and variance of the richness of a sample can be expressed using the expressions (1) replacing $p_i$ by $q_i$. The implication is that in the context of object-specific acceptance probabilities, richness depends only on the overall fraction of objects accepted and on the frequencies of objects among those accepted. When applied to genome editing libraries, richness depends only on the fraction of cells that are edited and on the frequencies of the edits within the edited cells. Frequencies of design reagents and design-specific probabilities of conferring edits are just intermediates and what matters for richness is their product.

### 2 Generalization of Richness to Require a Minimum Number of Observations

The concept of richness can be generalized to require more than just one observation of an object type. Define $R_{m,n}$ to be the number of object types with at least $n$ accepted counts in a random sample of size $m$.

$$\begin{aligned}
\mu_{m,n} =\ & S - \sum_{i=1}^{S} \sum_{k=0}^{n-1} \binom{m}{k} (fp_i)^k (1 - fp_i)^{m-k} \\
\sigma^2_{m,n} =\ & (2S-1)\mu_{m,n} - \mu^2_{m,n} - S(S-1) \\
& + \sum_{i=1}^{S} \sum_{\substack{j=1 \\ j \neq i}}^{S} \sum_{k=0}^{n-1} \sum_{l=0}^{n-1} \binom{m}{k} \binom{m-k}{l} (fp_i)^k (fp_j)^l (1 - fp_i - fp_j)^{m-k-l}
\end{aligned} \tag{2}$$

The derivation is provided in section 10.2

### 3 Approximations for Expected Value and Variance of Richness

If $m$ is large relative to $n$ the Poisson approximation to the binomial can be used to simplify. For the case $n = 1$ the approximation is guaranteed to be a lower bound for $\mu_{m,n}$ and an upper bound for $\sigma^2_{m,n}$.

$$\begin{aligned}
\lambda_i =\ & mfp_i \\
\gamma_i =\ & \sum_{k=0}^{n-1} \frac{\lambda_i^k e^{-\lambda_i}}{k!} \\
\mu_{m,n} =\ & S - \sum_i \gamma_i \\
\sigma^2_{m,n} =\ & \sum_i \gamma_i - \sum_i \gamma_i^2
\end{aligned} \tag{3}$$

When S is large this approximation can be helpful for computing the variance as it changes the computation from $O(S^2 n^2)$ to $O(Sn)$.

#### 3.1 Maximum value for $\sigma^2_{m,n}$

The approximation for $\sigma^2_{m,n}$ can be rewritten as

$$\sigma^2_{m,n} = \sum_i \gamma_i (1 - \gamma_i) \tag{4}$$

Each summand is of the form $x(1-x)$ where $x$ is in the range $[0, 1]$, so the maximum possible value will be $1/4$, which happens if and only if $\gamma_i = 1/2$. Thus the maximum possible value for $\sigma^2_{m,n}$ is $S/4$, which will be attained when $\mu_{m,n}$ is $S/2$.

### 4 Maximizing Richness

Consider that situation where the goal is to maximize the number of object types seen at least once ($n = 1$). The approximation for $\mu_m$ can be used to illustrate four different strategies to maximize richness.

$$\mu_m = S - \sum_i e^{-mfp_i}$$

- One obvious way to maximize $\mu_m$ would be to increase $m$, the number of samples taken from the distribution. In practice there can be limitations to this approach, for example if there are cost constraints associated with sampling.
- Another way would be to increase the acceptance probability $f$, getting it as close to 1 as possible.
- In situations where $m$, $f$ and $S$ are all fixed, the only remaining thing that could be modified to maximize $\mu_m$ is the distribution of object frequencies $p_i$. The expression for richness will be maximized when the summation is minimized. The summands are convex functions of $p_i$, which can be seen by noting that the second derivative with respect to $p_i$ is nonnegative. The sum of a convex function applied to a probability simplex is minimized when the probabilities are all equal, therefore $\mu_m$ is maximized when $p_i = 1/S$ and the maximum value is

$$\mu_m^{\texttt{max}} = S(1 - e^{-mf/S}) \tag{5}$$

- The final option is to increase $S$, the size of the library being screened. It won't always be possible - there may be contextual factors that put a limit on $S$. Increasing $S$ by a finite amount is not guaranteed to help - if a larger library has a less uniform frequency distribution it might actually result in lower richness compared to a smaller library. However it can be shown that in the limit, increasing $S$ far enough will help regardless of the library distribution.

  To see this note that $\mu_m$ can be written as $S\sum_i(1 - e^{-mfp_i})$. Using the Taylor series expansion for $e^x$ this can be rewritten as

$$\mu_m = S\left(\sum_i mfp_i - \frac{(mfp_i)^2}{2!} + \frac{(mfp_i)^3}{3!} + \dots\right)$$

  As the value of $S$ increases the values of $p_i$ will, on average, decrease, and the terms with larger powers of $p_i$ will decrease. For sufficiently large values of $S$ the first term will dominate and $\mu_m$ will tend towards $mfS$ where $f = \sum_i fp_i$.

### 5 Rule of Thumb for Richness

There is a relationship between the expected richness and the fraction of library members whose frequency exceeds a certain threshold. This can be useful as a rough guide for understanding richness.

Define $F_N$ to be the fractional richness of an $N$-fold sampling from a library of size $S$. In other words, $F_N$ is the number of unique types seen in a sample of size $NS$.

$$\begin{aligned} F_N =& \frac{1}{S} R_{NS,1} \\ E[F_N] =& \frac{1}{S} E[R_{NS,1}] \\ \approx& \frac{1}{S} \sum_i 1 - e^{-NSfp_i} \end{aligned} \tag{6}$$

Consider the terms in the sum, each representing the probability that type $i$ is not accepted among the $NS$ samples. Note that each term in the sum is of the form $1 - e^{-x}$. When $p_i = 0$ the summand evaluates to 0. As $p_i$ increases the summand increases from 0 to 1. The rate of transition is $NSf$ so as long as $f$ is not very small, the rate of transition will be very fast, making it effectively a phase transition. The phase transition from 0 to 1 will happen around the time when the term in the exponent is equal to $-1$, or equivalently around the point at which $p_i = \frac{1}{NSf}$.

In other words, the value of the sum will be very close to the number of types whose frequency is above $\frac{1}{NSf}$. This provides for a simple rule-of-thumb for estimating the expected richness. As an example, in the case that $f = 1$, the expected richness will be approximately equal to the number of types whose frequency in the original library exceeds $\frac{1}{NS}$.

### 6 Parametric Frequency Distributions

The relationships so far have been derived for general situations where the frequencies of different member types in a library can follow any distribution. In practical applications it is sometimes the case that frequency distributions can be well approximated by parametric distributions.

A natural distribution to consiser, and one that is often a good empirical fit, is the beta distribution. The beta distribution is commonly parameterized by shape parameters $\alpha$ and $\beta$ but for the current application it can be more convenient to parameterize the distribution by its mean and coeffieient of variation (CV). Since the distribution is being used to model $S$ probabilities which should sum to one, it makes sense to constrain the mean to be equal to $1/S$, leaving the coefficient of variation $c$ as the single free parameter. The usual beta shape parameters can be defined in terms of $S$ and $c$ as follows

$$\begin{aligned} \alpha =& \frac{S-1}{Sc^2} - \frac{1}{S} \\ \beta =& (S-1)\alpha \end{aligned} \tag{7}$$

If library member frequencies follow a beta distribution with a CV of $c$, the expected fractional richness associated with a sample of size $m$ from a library of size $S$ and with acceptance probability f, is given by

$$E\left[\frac{R_m}{S}\right] = 1 - (1 + Fc^2)^{-\left(\frac{1}{c^2}\right)} \tag{8}$$

where the fractional sampling $F$ is defined as $F = mf/S$. The expression (8) can be used to show that $\lim_{c^2 \to 0} E\left[\frac{R_m}{S}\right] = 1 - e^{-F}$, which is consistent with the upper bound described in expression (5).

For the more general situation where richness is based on needing to have at least $n$ accepted members of each type, the expected richness generalizes to

$$E\left[\frac{R_{m,n}}{S}\right] = \quad 1 - \sum_{k=0}^{n-1} \left(\frac{1}{1+Fc^2}\right)^{\frac{1}{c^2}} \left(1 - \frac{1}{1+Fc^2}\right)^k \binom{1/c^2 + k - 1}{k} \tag{9}$$

The derivation is provided in section 10.4

$$\begin{aligned} Var\left[\frac{R_{m,n}}{S}\right] = \quad & \frac{1}{S}\sum_{k=0}^{n-1} \left(\frac{1}{1+Fc^2}\right)^{\frac{1}{c^2}} \left(\frac{Fc^2}{1+Fc^2}\right)^k \binom{\frac{1}{c^2} + k - 1}{k} \\ & - \frac{1}{S}\sum_{k=0}^{n-1} \left(\frac{1}{1+2Fc^2}\right)^{\frac{1}{c^2}} \left(\frac{Fc^2}{1+2Fc^2}\right)^{2k} \binom{1/c^2 + k - 1}{k} \binom{1/c^2 + 2k - 1}{k} \end{aligned} \tag{10}$$

The derivation is provided in section 10.5

### 7 Maximizing Richness by Splitting Libraries

In some cases richness can be increased by splitting a library up into sub-libraries. This applies when a library can be split into sub-libraries where the CVs of frequencies in the sub-libraries are smaller than the CV in he original library. An example would be a library with a bi-modal frequency distribution, in which case splitting into two libraries corresponding to the two modes would result in sub-libraries each with lower CV than the original. This section considers the situation where richness is defined as the number of library members accepted at least once.

Start with a library of size $2S$ which can be split into two sub-libraries of size $S$ each. Let the type frequencies in the combined library be represented by $p_{a,i}$ and $p_{b,i}$ for $i = 1 \ldots S$, each with corresponding acceptance probabilities $f_a$ and $f_b$.

First consider the situation where the two sub-libraries have identical distributions, so $p_{a,i} = p_{b,i}$ and $f_a = f_b$ for all values of $i$. The expected richness for a sample of size $m$ from the combined library can then be expressed as

$$\mu_m = 2S - 2\sum_i e^{-mf_a p_{a,i}}$$

A natural stratified sampling approach would involve taking a sample of size $m/2$ from each of the sub-libraries $A$ and $B$. Within each sub-library the frequencies will be double what they were in the original library, therefore the expected richness from each sub-library will be $S - \sum_i e^{-\frac{m}{2}2f_a p_{a,i}}$ which is exactly half the expected richness for a sample of size $m$ from the original library. In other words, the stratified sampling approach provides no improvement in expected richness.

Now consider a library of size S which can be split into two sub-libraries $A$ and $B$ of size $S_a$ and $S_b$ where the product of frequency and acceptance probability is identical within each sub-library - so $fp_i = f/S_a$ for all members of sub-library $A$ and $fp_i = f/S_b$ for all members of sub-library $B$. If the total sample size is $m$ then a natural way to stratify is to to sample $mS_a/S$ members from library $A$ and $mS_b/S$ from library $B$. As a result, the product of the sample size, frequency and acceptance probability in the $\lambda_i$ terms in equation 3 are all equal to $mf/S$ and the expected richness will attain the maximum value described in equation 5.

Another extreme to consider is to split the library into $m$ sub-libraries each of size 1, sampling once from each (it is assumed that $m \leq S$). In this situation the richness is guaranteed to take on the maximum possible value of $m$.

The general rule is that whenever it is possible and cost-effective to stratify a library into sub-libraries within which the CV of frequencies is lower, it will improve richness. Conversely, if the CV of frequencies in the proposed strata is the same as in the combined library, a stratified sampling approach will provide no improvement in richness.

Figure 1 provides an example of a library with some opportunity to improve richness via a stratified sampling approach, due to having a bi-modal frequency distribution.

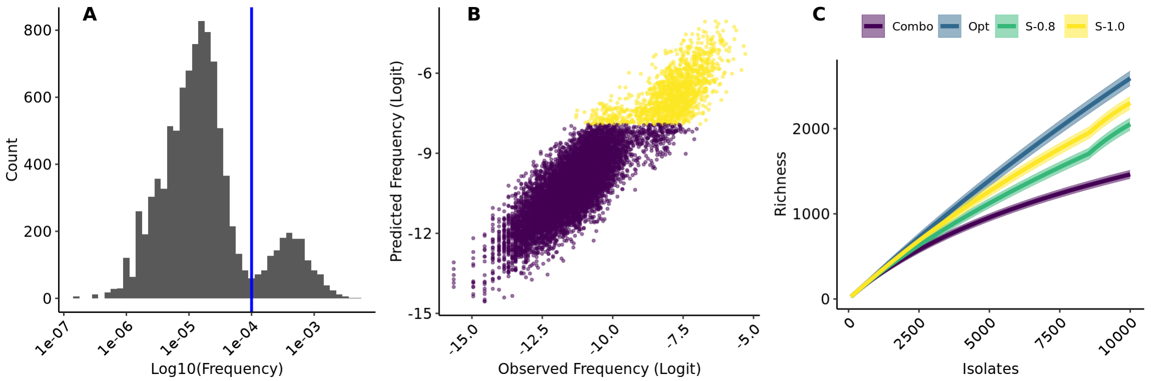


Figure 1: Example of the benefit of splitting a library. **A**: Histogram of design frequencies for a library targeting 10002 edits. The design frequencies follow a sub-optimal bi-modal distribution where 15% of the designs have a frequency roughly 30 times larger than the remaining 85%. **B**: Comparison between empirically-determined design frequencies and predictions from a model using design sequence properties as input. The $R^2$ of the predictive model is 0.8 and the color of the points indicates a partition of the designs into two classes by clustering with a Gaussian mixture model fit on the predictions. **C**: Richness in a screen of 10,000 isolates, assuming an edit efficiency of 30% for all designs. The red 'Combo' curve shows the expected richness when screening with all designs combined into a single library. The green 'Opt' curve is the theoretical optimum for a perfectly even library where all edits are present at the same frequency. The blue 'S-0.8' curve shows predicted richness when splitting the designs into two libraries using a Gaussian mixture model fitted to the predicted frequencies, as indicated in panel B. The number of isolates screened from each library is proportional to the library size, the discontinuity in the richness curve is the point of transition from screening one library to the other. The magenta 'S-1.0' curve is as in the previous curve but splitting into two libraries based on predictions that are perfectly correlated with the empirical frequencies

### 8 Predicting Sampling Depth Requirements

Parametric distributions of library member frequencies can be useful to model scenarios prior to generating experimental data. As an example, the beta model can be used to predict the sampling depth that would be required to observe a certain fraction of the members in the library.

Consider a genome engineering library in which the edit frequencies follow a beta distribution. If the edit frequencies are to be estimated by sequencing the DNA of a sample of cells from the library, it is helpful to think about the relationship between the number of cells that will be sampled and the confidence interval for the estimate of each edit's frequency.

For an edit present at frequency $p$, taking a sample of $10/p$ cells will result in a 95% confidence interval of $(0.4p, 1.7p)$, so the estimate for the edit's frequency will be accurate to within roughly 60-70% of the true value. Thus setting the sampling depth to $10/p$ can be used as an approximate guide to thinking about how deeply to sample.

For example, in a 1000-member edit library with a CV of 2, the 10th percentile of the edit frequency distribution is $2.6 \times 10^{-7}$ (Figure 2). Using the above approximate rule, one would want a sampling depth of around $2.6 \times 10^{8}$ to get an approximate estimate of frequency for 90% of the library. In the context of profiling and *E. coli* library (with a 4.6Mb genome) using 2x150 short read sequencing, a total of $4 \times 10^{12}$ read pairs would be required. If the fraction of cells that are edited is less than 1, the estimate would need to be inflated accordingly.

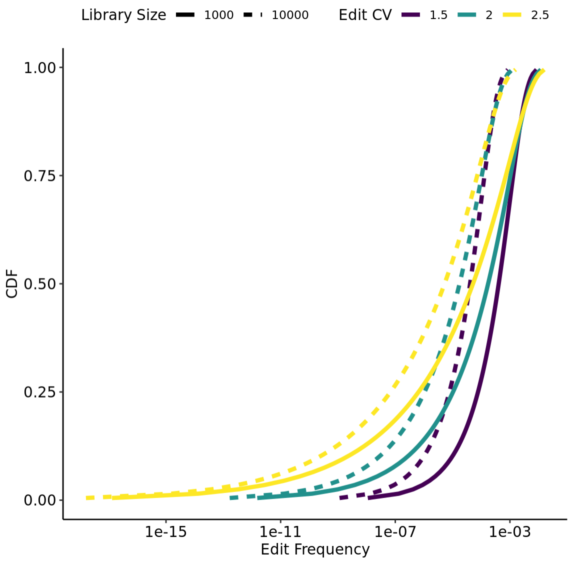


Figure 2: Cumulative distribution function for edit frequency at a variety of library sizes and CV, assuming a beta distribution for edit frequency.

### 9 Estimation of CV for a Frequency Distribution

Section 6 indicates that CV is a critical statistic influencing richness outcomes. This section considers the problem of estimating CV for a frequency distribution.

Consider the problem of estimating the CV for a distribution of frequencies for a library of size S. Many practical applications can be modeled as a compound binomial process where frequencies are randomly generated from an underlying distribution with mean $1/S$ and where the number of observations of each library member is a binomial random variable.

In a genome editing example, a common situation would be use of short read sequencing in regions targeted for editing to determine the frequencies of each of the edits. The underlying edit frequencies follow some distribution, and for each edit the number of sequence reads corresponding to the edit would be a binomial variable.

Let the underlying edit frequencies follow an arbitrary distribution with probability density function $f(p)$ for $p \in [0, 1]$. Let $X_i$ be the number times member $i$ is observed among $n$ samples. In the edit detection context, $X_i$ would be the number of sequencing reads classified as supporting edit $i$ and $n$ would be the total read depth at the targeted genomic site. This will include reads from DNA of cells in the edited cell library that are targeted to have edit $i$, in a background of DNA from cells in the library that are targeted to have other edits.

The probability that random variable $X_i$ takes on value $k$ is given by

$$P(X_i = k) = \int_0^1 \binom{n}{k} p^k (1-p)^{n-k} f(p) dp$$

#### 9.1 CV for a Compound Binomial Model

The parameter of key importance for predicting richness is the CV $c$ of the underlying frequency distribution. Given $N$ binomial observations $X_i$ for $i = 1 \ldots N$ where each count results from $n$ trials, there are a few options for estimating the CV. Let $Z_i = X_i/n$ be the observed edit frequencies.

#### 9.2 Compound Binomial CV Estimation Using Sample Variance

One approach for estimation of CV is to use the usual sample variance $V$, defined as

$$V = \frac{1}{N-1}\sum_{i=1}^{N}\left(Z_i - \bar{Z}\right)^2$$
$$\bar{Z} = \frac{1}{N}\sum_{i=1}^{N} Z_i \tag{11}$$

A natural estimator for the CV is the ratio of the standard deviation of the $Z_i$ values to their mean, call this estimate $\hat{c_{\texttt{naive}}}$. Under the assumption that the mean of the frequency distribution is $1/S$, the estimator is defined as $\hat{c_{\texttt{naive}}} = S\sqrt{V}$. A problem with this intuitive approach is that it is positively biased, sometimes quite severely so. Section 10.6.2 shows that the bias of $\hat{c_{\texttt{naive}}}$ is given by

$$\texttt{Bias}[\hat{c_{\texttt{naive}}}] = \sqrt{\frac{S-1}{n} + \frac{(n-1)}{n}c^2} - c \tag{12}$$

Figure 3 presents the magnitude of bias in a few example scenarios.

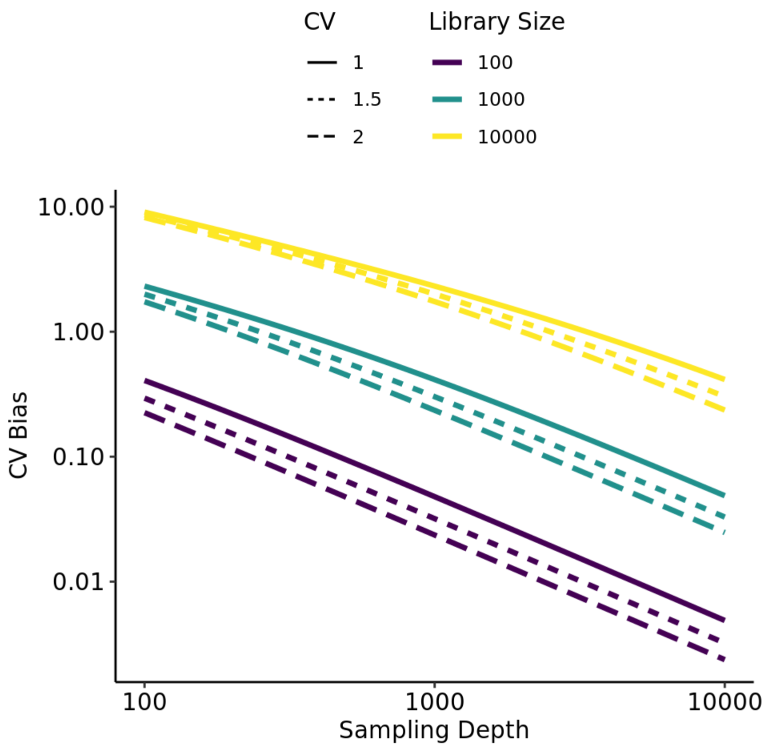


Figure 3: Bias of the CV estimator $\hat{c_{\texttt{naive}}}$ for a range of different conditions. The bias, described in equation (12), depends only on the CV of the underlying probability distribution, the size of the library, and the sampling depth. When library size is large relative to the sampling depth the size of the bias can be substantial. For example, if sampling 100 observations from a 10,000-edit library with a CV of 1, the bias of $\hat{c_{\texttt{naive}}}$ is 9.0 resulting in an estimate of CV that is around 10-fold larger than the true value.

This motivates a modified estimator $\hat{c^2}$ based on the sample variance $V$, with terms to correct for the above bias

$$\hat{c^2} = \frac{n}{n-1} S^2 V - \frac{S-1}{n-1} \tag{13}$$

Note that $\hat{c^2}$ is not guaranteed to be positive. In cases where $S$ is large relative to $n$ and/or $c$ is large, the sample variance can be small relative to the quantity subtracted from it, resulting in $\hat{c^2}$ becoming negative. Nevertheless, $\hat{c^2}$ is an unbiased estimator of $c^2$.

In situations where $\hat{c^2}$ is used, it can be helpful to know its variance, which can be expressed as

$$Var[\hat{c^2}] = \left(\frac{n}{n-1}\right)^2 S^4 \left(\frac{\mu_4^c}{N} - \frac{(\mu_2^c)^2(N-3)}{N(N-1)}\right) \tag{14}$$

Where $\mu_j^c$ is the nth central moment of the random variable $Z$. Use of the delta method demonstrates that

$$Var\left[\sqrt{\hat{c^2}}\right] = \left(\frac{n}{n-1}\right)^2 \frac{S^4}{4c^2} \left(\frac{\mu_4^c}{N} - \frac{(\mu_2^c)^2(N-3)}{N(N-1)}\right) \tag{15}$$

Derivations of the above expressions are presented in section 10.6

#### 9.3 Compound Binomial CV Estimation Using Maximum Likelihood

The CV estimation methods introduced so far make no assumption on the distributional form of the edit frequencies. Another standard approach to estimate CV is to assume a parametric distribution for $f()$ and to use maximum likelihood estimation (MLE) to estimate the CV. If $f()$ is assumed to be a beta distribution, the counts $X_i$ follow the well-studied beta-binomial distribution for which MLE is straightforward.

Figure 4 evaluates the performance of a parametric fit using a beta binomial model, comparing it with other methods that have recently been used for predicting richness in sequencing libraries (Daley and Smith 2013). In the figure, two edit libraries are studied: a 10,000-design E. coli library and a 6,000-design yeast library. A single whole genome shotgun library is made from each pool of edited cells and both sequencing libraries are sequenced together on two NextSeq 2x150 runs.

For each organism, one sequencing run is used as a training dataset to predict the Edit Fractional Richness that will be observed in the other sequencing run. To explore the performance of predictions as a function of sample size, the training dataset is randomly downsampled to differing levels of genome coverage, as measured by the median coverage depth across the targeted edit regions. For each coverage depth, 1000 randomly downsampled datasets are made and each is used to train a richness predictor which is then used to predict edit fractional richness at the sampling depth encountered in the withheld test dataset.

The performance of the predictors is summarized as the root mean squared error (RMSE) where error is defined as the difference between the predicted and observed fractional richness in the test data. The dashed lines span the inter-quartile range of RMSE. The fractional richness predictions depend on the Edit Fraction, which is estimated as the sum of the individual edit frequencies (the count of reads in which an edit is found divided by the total genomic depth at the edited locus).

Three predictors of richness are evaluated. “BB” fits a beta-binomial model using maximum likelihood estimation to fit a single parameter, the CV, then uses relationship (8) to predict fractional richness in the test data. The size of the library is assumed to be known, which is typically the case for edited cell libraries. “DS” is the nonparametric Daley-Smith estimator (Daley and Smith 2013) and “ZTNB” is a parametric approach used in the same body of work to fit a zero-truncated negative binomial model. Neither the DS nor the ZTNB approach makes an assumption about the size of the design library, it is inferred from the training data.

At lower sampling depths the BB model performs better, likely due to the fact that the edit frequencies are well described by a beta binomial model, thus the model can be leveraged to make good predictions from limited data. At higher coverage the gap between the methods mostly narrows. An interesting exception is the E. coli run2 dataset, which has a heavier left tail with some designs dropping out. The DS and ZTNB models don't assume the library size, which leads to less accurate prediction when sample size is small and not much information is available to estimate the library size, but with deep sampling these approaches are able to learn that the effective library size is lower than ideal, whereas the BB approach assumes a fixed library size and thus over-estimates the fractional richness.

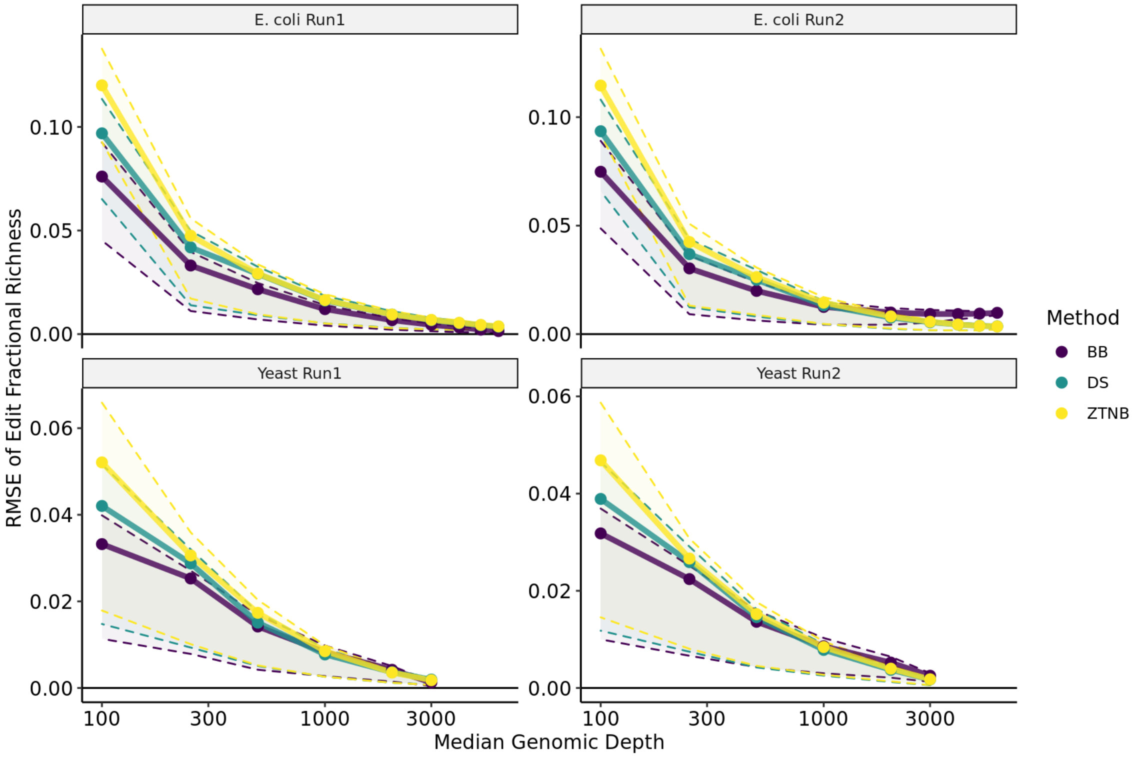


Figure 4: Three predictors of richness are compared. "BB" fits a beta-binomial model using maximum likelihood to estimate a single parameter, the CV, then uses relationship (8) to predict fractional richness in a separate test data set. "DS" is the nonparametric Daley-Smith estimator (Daley and Smith 2013) and "ZTNB" is a parametric approach used in the same body of work to fit a zero-truncated negative binomial model. Unlike the BB approach, neither the DS nor the ZTNB approach makes an assumption about the size of the design library; library size is inferred from the training data. At lower sampling depths the BB model performs better, likely due to the fact that the edit frequencies are well described by a beta binomial model, hence the assumed model can be leveraged to make good predictions from limited data. However, with deep sampling the DS and ZTNB approaches have lower error than the BB approach, likely due to having more flexibility to adapt to the observed data in the context of a large sample size.

### 10 Detailed Derivations

#### 10.1 Derivation of Exact Expressions for $\mu_m$ and $\sigma_m^2$

$R_m$, the richness of a sample of size $m$, can be defined as $\sum_i 1_i$ where $1_i$ is an indicator variable with value 1 if a member type $i$ is sampled and accepted at least once among the $m$ items sampled, and is 0 otherwise.

Note that $E[1_i] = P(\text{type } i \text{ is accepted at least once})$ and $E[1_i^2] = E[1_i]$, relationships that will be used during the derivation.

$\mu_m$ can be written as

$$\begin{aligned}\mu_m =& E\left[\sum_i 1_i\right] \\ =& \sum_i E[1_i] \\ =& \sum_i P(\text{type } i \text{ accepted at least once among the } m \text{ samples}) \\ =& \sum_i \left(1 - P(\text{type } i \text{ not accepted among the } m \text{ samples})\right)\end{aligned} \tag{16}$$

Type $i$ will be absent in a sample if either type $i$ is not selected, or if it is selected and not accepted. Thus $P(\text{type } i \text{ not accepted in one sample})$ is equal to $(1-p_i)+p_i(1-f)$ which simplifies to $1-fp_i$. That means $P(\text{type } i \text{ not accepted among the } m \text{ samples}) = (1-fp_i)^m$. Putting this back into (16):

$$\begin{aligned}\mu_m =& \sum_i 1-(1-fp_i)^m \\ =& S - \sum_i (1-fp_i)^m\end{aligned} \tag{17}$$

To compute the variance of $R_m$ we use the relationship $Var[X] = E[X^2] - (E[X])^2$.

$$\begin{aligned}\sigma_m^2 =& E[R_m^2] - \mu_m^2 \\ E[R_m^2] =& E\left(\sum_i \sum_j 1_i 1_j\right) \\ =& \sum_i \sum_j E[1_i 1_j] \\ =& \sum_i E[1_i^2] + \sum_i \sum_{j\neq i} E[1_i 1_j] \\ =& \sum_i E[1_i] + \sum_i \sum_{j\neq i} P(\text{type } i \text{ and } j \text{ both accepted}) \\ =& \mu_m + \sum_i \sum_{j\neq i} P(\text{type } i \text{ and } j \text{ both accepted})\end{aligned} \tag{18}$$

For $P(\text{type } i \text{ and } j \text{ both accepted among } m \text{ samples})$ we can use the relationship

$$P(A\&B) = 1 - P(\sim A) - P(\sim B) + P(\sim (A|B))$$

where ∼, & and | indicate the logic operations NOT, AND and OR respectively. Using this relationship we can express $P$(type $i$ and $j$ are both accepted among $m$ samples) as

$$1 - P(i \text{ not accepted}) - P(j \text{ not accepted}) + P(\text{neither } i \text{ nor } j \text{ accepted})$$

The probability that type $i$ is not accepted among the $m$ samples was shown above to be $(1-fp_i)^m$. So the only thing left to compute is the probability that neither type $i$ nor type $j$ is accepted among the m samples. First consider taking just one sample - if neither type $i$ nor type $j$ is accepted then it means one of three things has happend:

- Neither type $i$ nor type $j$ is sampled, which happens with probability $(1 - p_i - p_j)$
- Type $i$ is sampled and not accepted, which happens with probability $p_i(1 - f)$
- Type $j$ is sampled and not accepted, which happens with probability $p_j(1 - f)$

Summing the probability of these three disjoint events simplifies to $1 - fp_i - fp_j$, and to get the probability that neither type $i$ nor type $j$ is accepted among all $m$ samples the term is raised to the power $m$.

Putting this back into (18) we get

$$\begin{aligned} E[R_m^2] &= \mu_m + \sum_i \sum_{j \neq i} (1 - (1 - fp_i)^m - (1 - fp_j)^m + (1 - fp_i - fp_j)^m) \\ Var[R_m] &= E[R_m^2] - \mu_m^2 \\ &= \mu_m - \mu_m^2 + \sum_i \sum_{j \neq i} (1 - (1 - fp_i)^m - (1 - fp_j)^m + (1 - fp_i - fp_j)^m) \end{aligned} \tag{19}$$

The first term in the double summation is constant and thus can be pulled out and expressed as $S(S-1)$.

The second and third terms in the double summation iterate over just one of the variables and can each be rewritten as $(S-1)\sum_i (1 - fp_i)^m$ and noting the similarity with the formula for $\mu_m$ the sum of the second and third terms is equal to $2(S-1)(\mu_m - S)$. Then, after a little rearrangement, the expression for variance simplifies to

$$Var[R_m] = (2S-1)\mu_m - \mu_m^2 - S(S-1) + \sum_i \sum_{j \neq i} (1 - fp_i - fp_j)^m \tag{20}$$

#### 10.2 Derivation of Exact Expressions for $\mu_{m,n}$ and $\sigma^2_{m,n}$

$R_{m,n}$, the number of library members seen at least $n$ times in a sample of size $m$, can be defined as $\sum_i 1_i$ where $1_i$ is an indicator variable with value 1 if at least $n$ members of type $i$ are accepted among the $m$ items sampled, and is 0 otherwise.

$\mu_{m,n}$ can be written as

$$\begin{aligned} \mu_{m,n} &= E\left[\sum_i 1_i\right] \\ &= \sum_i E[1_i] \\ &= \sum_i P(\text{type } i \text{ accepted at least } n \text{ times among } m \text{ samples}) \\ &= \sum_i (1 - P(\text{type } i \text{ accepted less than } n \text{ times among } m \text{ samples})) \end{aligned} \tag{21}$$

The number of members of type $i$ accepted is a binomial random variable with success probability $fp_i$, so the probability of having fewer than $n$ among $m$ samples is given by the binomial tail probability $\sum_{k=0}^{n-1} \binom{m}{k} (fp_i)^k (1 - fp_i)^{m-k}$

Putting this back into (21):

$$\mu_{m,n} = S - \sum_{i=1}^{S} \sum_{k=0}^{n-1} \binom{m}{k} (fp_i)^k (1 - fp_i)^{m-k} \tag{22}$$

To compute the variance of $R_{m,n}$ we use the relationship $Var[X] = E[X^2] - (E[X])^2$.

$$\begin{aligned}
\sigma^2_{m,n} =& E[R^2_{m,n}] - \mu^2_{m,n} \\
E[R^2_{m,n}] =& E\left[\sum_i \sum_j 1_i 1_j\right] \\
=& \sum_i \sum_j E[1_i 1_j] \\
=& \sum_i E[1_i^2] + \sum_i \sum_{j \neq i} E(1_i 1_j) \\
=& \sum_i E[1_i] + \sum_i \sum_{j \neq i} P(\text{types } i \text{ and } j \text{ both accepted at least } n \text{ times}) \\
=& \mu_{m,n} + \sum_i \sum_{j \neq i} P(\text{types } i \text{ and } j \text{ both accepted at least } n \text{ times})
\end{aligned} \tag{23}$$

For $P(\text{types } i \text{ and } j \text{ both accepted at least } n \text{ times})$ we use the relationship

$$P(A \& B) = 1 - P(\sim A) - P(\sim B) + P(\sim (A|B))$$

to express it as

$$1 - P(i \text{ accepted less than } n \text{ times}) - P(j \text{ accepted less than } n \text{ times}) + P(i \text{ and } j \text{ each accepted less than } n \text{ times})$$

The probability that member type $i$ is accepted fewer than $n$ times has been derived above. The probability that both type $i$ and type $j$ are each accepted less than $n$ times can be written as a sum of multinomial probabilities $\sum_{k=0}^{n-1} \sum_{l=0}^{n-1} \binom{m}{k} \binom{m-k}{l} (fp_i)^k (fp_j)^l (1 - fp_i - fp_j)^{m-k-l}$

Putting this back into (23) we get

$$
\begin{aligned}
E[R^2_{m,n}] = \quad & \mu_{m,n} + \sum_i \sum_{j \neq i} \Bigg[ 1 - \sum_{k=0}^{n-1} \binom{m}{k} (fp_i)^k (1-fp_i)^{m-k} - \sum_{k=0}^{n-1} \binom{m}{k} (fp_j)^k (1-fp_j)^{m-k} \\
& + \sum_{k=0}^{n-1} \sum_{l=0}^{n-1} \binom{m}{k} \binom{m-k}{l} (fp_i)^k (fp_j)^l (1-fp_i-fp_j)^{m-k-l} \Bigg] \\
= \quad & \mu_{m,n} + S(S-1) - 2(S-1)(S-\mu_{m,n}) \\
& + \sum_i \sum_{j \neq i} \sum_{k=0}^{n-1} \sum_{l=0}^{n-1} \binom{m}{k} \binom{m-k}{l} (fp_i)^k (fp_j)^l (1-fp_i-fp_j)^{m-k-l} \\
= \quad & (2S-1)\mu_{m,n} - S(S-1) \\
& + \sum_i \sum_{j \neq i} \sum_{k=0}^{n-1} \sum_{l=0}^{n-1} \binom{m}{k} \binom{m-k}{l} (fp_i)^k (fp_j)^l (1-fp_i-fp_j)^{m-k-l} \\
\sigma^2_{m,n} = & E[R^2_{m,n}] - \mu^2_{m,n} \\
= \quad & (2S-1)\mu_{m,n} - \mu^2_{m,n} - S(S-1) \\
& + \sum_i \sum_{j \neq i} \sum_{k=0}^{n-1} \sum_{l=0}^{n-1} \binom{m}{k} \binom{m-k}{l} (fp_i)^k (fp_j)^l (1-fp_i-fp_j)^{m-k-l}
\end{aligned}
\tag{24}
$$

#### 10.3 Derivation of Approximations for $\mu_{m,n}$ and $\sigma^2_{m,n}$

The expected value of richness was derived in (22)

$$
\mu_{m,n} = S - \sum_{i=1}^{S} \sum_{k=0}^{n-1} \binom{m}{k} (fp_i)^k (1-fp_i)^{m-k}
$$

Defining $\lambda_i = mfp_i$, the summands can be rewritten as

$$
\begin{aligned}
\binom{m}{k} (fp_i)^k (1-fp_i)^{m-k} = \quad & \frac{m(m-1)\dots(m-k+1)}{k!} \left(\frac{\lambda_i}{m}\right)^k \left(1 - \frac{\lambda_i}{m}\right)^{m-k} \\
\approx \quad & \frac{\lambda_i^k}{k!} \left(1 - \frac{\lambda_i}{m}\right)^m \quad \text{if } m \text{ is large relative to } k \\
\approx \quad & \frac{\lambda_i^k e^{-\lambda_i}}{k!} \quad \text{if } m \text{ is large}
\end{aligned}
\tag{25}
$$

The same kind of logic simplifies the summands in the expression for $\sigma^2_{m,n}$ in (24)

$$
\begin{aligned}
&\binom{m}{k}\binom{m-k}{l}(fp_i)^k(fp_j)^l(1-fp_i-fp_j)^{m-k-l} \\
= &\left(\frac{m\ldots(m-k+1)}{k!}\right)\left(\frac{(m-k)\ldots(m-k-l+1)}{l!}\right)\left(\frac{\lambda_i}{m}\right)^k\left(\frac{\lambda_j}{m}\right)^l\left(1-\frac{\lambda_i}{m}-\frac{\lambda_j}{m}\right)^{m-k-l} \\
\approx &\frac{\lambda_i^k}{k!}\frac{\lambda_j^l}{l!}\left(1-\frac{\lambda_i}{m}-\frac{\lambda_j}{m}\right)^m \quad \text{if } m \text{ is large relative to } k \text{ and } l \\
\approx &\frac{\lambda_i^k\lambda_j^l e^{-(\lambda_i+\lambda_j)}}{k!l!} \quad \text{if } m \text{ is large} \\
= &\frac{\lambda_i^k e^{-\lambda_i}}{k!}\frac{\lambda_j^l e^{-\lambda_j}}{l!}
\end{aligned} \tag{26}
$$

Plugging the Poisson approximation back in to (24) and rearranging leads to the simplified expression in (3)

#### 10.4 Derivation of Fractional Richness for a Beta Distribution

For a library of size $S$ with frequences following a beta distribution with CV of $c$, the mean will be $1/S$ and the distribution's commonly-used shape parameters $\alpha$ and $\beta$ are given by

$$
\begin{aligned}
\alpha =& \frac{S-1}{Sc^2} - \frac{1}{S} \\
\beta =& (S-1)\alpha
\end{aligned} \tag{27}
$$

Using the expression for richness in (3), the fractional richness can be expressed as

$$
\frac{\mu_{m,n}}{S} = 1 - \frac{1}{S}\sum_{i=1}^{S}\sum_{k=0}^{n-1}\frac{(mfp_i)^k e^{-mfp_i}}{k!}
$$

If the frequencies $p_i$ follow a gamma distribution with shape $\alpha$ and rate $\beta$, the fractional richness can be expressed as

$$
\begin{aligned}
\frac{\mu_{m,n}}{S} &= 1 - \int_0^1 \sum_{k=0}^{n-1}\frac{(mfp)^k e^{-mfp}}{k!}\frac{\Gamma(\alpha+\beta)}{\Gamma(\alpha)\Gamma(\beta)}p^{\alpha-1}(1-p)^{\beta-1}dp \\
&= 1 - \sum_{k=0}^{n-1}\frac{\Gamma(\alpha+\beta)}{\Gamma(\alpha)\Gamma(\beta)}\frac{(mf)^k}{k!}\int_0^1 p^{\alpha+k-1}(1-p)^{\beta-1}e^{-mfp}dp
\end{aligned}
$$

For large libraries the value of $S$ should greatly exceed the CV and the value of $\beta$ will be very large. In that case, two approximations can be used to make substitutions in the expression for fractional richness

$$
\begin{aligned}
\frac{\Gamma(\alpha+\beta)}{\Gamma(\beta)} &\approx \beta^\alpha \\
(1-p)^{\beta-1} &\approx e^{-(\beta-1)p}
\end{aligned}
$$

The accuracy of the first of the above two approximations is characterized in figure 5. The second approximation is based on the Maclaurin series for the exponential function which is very accurate when p is small, and when p is large the fact that $\beta$ is large means the term and thus the error will be small.

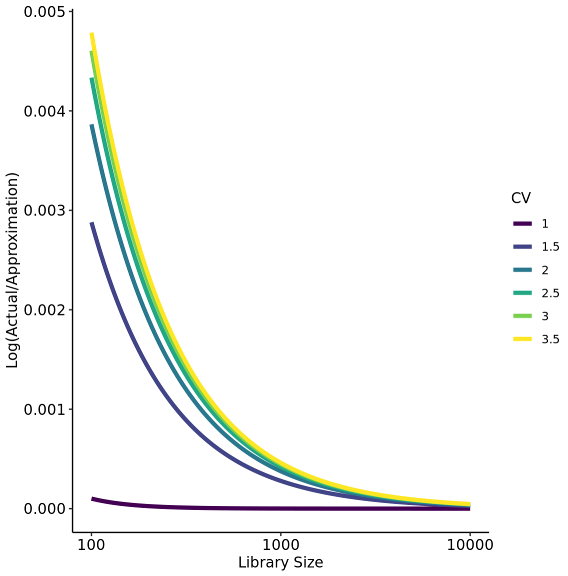


Figure 5: Assessment of accuracy of the approximation of $\Gamma(\alpha+\beta)/\Gamma(\beta)$ by $\beta^\alpha$, for a range of values relevant to study of edit libraries. The log ratio of the actual over the approximation is plotted against library size for a range of different CV values. Library size and CV determine the parameters $\alpha$ and $\beta$ as defined in (27). The error of the approximation is largest when library size is small and CV is large. The largest log ratio over the range of parameters evaluated is 0.0048, meaning the approximation is 0.48% larger than the actual value, resulting in the approximation for richness being slightly lower than the exact value.

$$\frac{\mu_{m,n}}{S} = 1 - \sum_{k=0}^{n-1} \frac{\beta^\alpha}{\Gamma(\alpha)} \frac{(mf)^k}{k!} \int_0^1 p^{\alpha+k-1} e^{-(\beta-1+mf)p} dp$$

Making the variable substitution $x = (\beta - 1 + mf)p$ turns the integral into a lower incomplete gamma function and when $\beta$ is large, the upper limit of the integral is large and it is well appoximated by the complete gamma function. For notational convenience the $\beta - 1$ terms in the denominator can be replaced by $\beta$.

$$\begin{aligned}\frac{\mu_{m,n}}{S} &= 1 - \sum_{k=0}^{n-1} \left(\frac{\beta}{\beta-1+mf}\right)^\alpha \left(\frac{mf}{\beta-1+mf}\right)^k \frac{1}{\Gamma(\alpha)k!} \int_0^{\beta-1+mf} x^{\alpha+k-1} e^{-x} dx \\ &= 1 - \sum_{k=0}^{n-1} \left(\frac{\beta}{\beta+mf}\right)^\alpha \left(\frac{mf}{\beta+mf}\right)^k \frac{\Gamma(\alpha+k)}{\Gamma(\alpha)k!}\end{aligned}$$

The parameters $\alpha$ and $\beta$ are defined in terms of the library size and frequency CV as in (7). For large values of $S$ they are well approximated by the simpler expressions $\alpha \approx 1/c^2$ and $\beta \approx S/c^2$, and the expression for fractional richness simplifies

$$\frac{\mu_{m,n}}{S} = 1 - \sum_{k=0}^{n-1} \left(\frac{1}{1+Fc^2}\right)^{\frac{1}{c^2}} \left(1 - \frac{1}{1+Fc^2}\right)^k \binom{1/c^2+k-1}{k} \tag{28}$$

The terms in the summation are of the form of a negative binomial distribution with size $1/c^2$ and probability $1/(1+Fc^2)$ which makes for very straightforward computational evaluation. It follows that the number of observations of each edit in the sample is distributed as a negative binomial.

This result is analogous to the fact that a Poisson-gamma mixture is distributed as a negative binomial distribution, noting that for large values of $\beta$ the beta distribution is well approximated by a gamma. In this context, a library member's frequency $p$ is sampled from a gamma distribution and the number of occurences in a sample of size $m$ and with acceptance rate $f$ is distributed as a Poisson random variable with rate $mfp$, with the effect that the distribution of the final accepted count follows a negative binomial distribution

#### 10.5 Variance of Fractional Richness for a Beta Distribution

The expression for variance of richness (3) can be used to determine the variance of fractional richness under the assumption that the probabilities follow a beta distribution

$$\frac{\sigma^2_{m,n}}{S} = \frac{1}{S}\sum_{i=1}^{S}\sum_{k=0}^{n-1}\frac{(mfp_i)^k e^{-mfp_i}}{k!} - \frac{1}{S}\sum_{i=1}^{S}\sum_{k=0}^{n-1}\left(\frac{(mfp_i)^k e^{-mfp_i}}{k!}\right)^2$$

Following the same kind of process as described in the previous section, it can be shown that

$$\begin{aligned}\frac{\sigma^2_{m,n}}{S} = {} & \frac{1}{S}\sum_{k=0}^{n-1}\left(\frac{1}{1+Fc^2}\right)^{\frac{1}{c^2}}\left(\frac{Fc^2}{1+Fc^2}\right)^k\binom{\frac{1}{c^2}+k-1}{k} \\ & - \frac{1}{S}\sum_{k=0}^{n-1}\left(\frac{1}{1+2Fc^2}\right)^{\frac{1}{c^2}}\left(\frac{Fc^2}{1+2Fc^2}\right)^{2k}\binom{1/c^2+k-1}{k}\binom{1/c^2+2k-1}{k}\end{aligned} \tag{29}$$

#### 10.6 Properties of CV Estimator

This section expands on the details underlying the estimator introduced in section 9.

For $i = 1 \ldots N$, define $X_i$ to be independent compound-binomial random variables with probabilities $P_i$ drawn from a distribution with density function $f()$ and with binomial size parameter $n$. Let $\mu^c_j$ be the $j$th central moment for $X_i$. Then $\hat{c^2}$ as defined in (13) is an unbiased estimate of $c^2$ where $c$ is the CV of $f()$, and its variance is as defined in (14).

This is demonstrated in three steps.

- In section 10.6.1, the variance of the sample variance estimator is derived in terms of $\mu^c_j$
- In section 10.6.2, $\mu^c_j$ is expressed in terms of the raw moments $\nu_j$ of the underlying probability $P$
- In section 10.6.3, estimation of the raw moments $\nu_j$ is addressed

###### 10.6.1 Variance of the Sample Variance

Given independent and identically distributed random variables $Z_i$ for $i = 1 \ldots N$, the expression for the sample variance $V$ can be written as

$$V = \binom{N}{2}^{-1} \sum_{i=1}^{N-1} \sum_{j=i+1}^{N} \frac{1}{2} (Z_i - Z_j)^2 \tag{30}$$

$V$ is an unbiased estimator for variance because $E[(Z_i - Z_j)^2/2] = \mu_2^c$ for all $i, j$.

$$\begin{aligned} Var[V] =& E[V^2] - (\mu_2^c)^2 \\ =& \binom{N}{2}^{-2} E\left[\sum_{i=1}^{N-1} \sum_{j=i+1}^{N} \left(\frac{1}{2} (Z_i - Z_j)^2 - (\mu_2^c)^2\right)\right]^2 \end{aligned} \tag{31}$$

The square of the summation involves terms of the following form

$$\left(\frac{1}{2} (Z_i - Z_j)^2 - (\mu_2^c)^2\right)\left(\frac{1}{2} (Z_k - Z_l)^2 - (\mu_2^c)^2\right) \tag{32}$$

There are three possible values for the expected value of each of the terms.

- if $i \neq k$ and $j \neq l$ the expected value is zero
- if $i = k$ and $j = l$ the expected value is $(\mu_4 + (\mu_2^c)^2)/2$. There are $\binom{N}{2}$ such terms
- otherwise the expected value is $(\mu_4 - (\mu_2^c)^2)/4$ and there are $N(N-1)(N-2)$ such terms

As a result it follows that

$$Var[V] = \frac{\mu_4^c}{N} - \frac{(\mu_2^c)^2(N-3)}{N(N-1)} \tag{33}$$

###### 10.6.2 Moments of a Compound-Binomial Distribution

Define $\mu_j^r$ to be the $j$th raw moment of the distribution of $Z$, $E[Z^j]$.

$$\begin{aligned} \mu_j^r =& \sum_{k=0}^{n} \frac{k^j}{n^j} \int_0^1 \binom{n}{k} p^k (1-p)^{n-k} f(p) dp \\ =& \int_0^1 \frac{1}{n^j} \left(\sum_{k=0}^{n} k^j \binom{n}{k} p^k (1-p)^{n-k}\right) f(p) dp \end{aligned} \tag{34}$$

The terms in parentheses are the raw moments for the binomial distribution. If $B$ is a random binomial variable with probability $p$ and size $n$, the first four raw moments are

$$\begin{aligned} E[B] =& np \\ E[B^2] =& np + n(n-1)p^2 \\ E[B^3] =& np + 3n(n-1)p^2 + n(n-1)(n-2)p^3 \\ E[B^4] =& np + 7n(n-1)p^2 + 6n(n-1)(n-2)p^3 + n(n-1)(n-2)(n-3)p^4 \end{aligned} \tag{35}$$

Replacing the inner sum by the appropriate expression for binomial moments, the moments of $Z$ can be expressed as linear combinations of the moments of the underlying frequency $P$, defined as $\nu_j = E[P^j]$

$$
\begin{aligned}
\mu_1^r =& \nu_1 \\
\mu_2^r =& \frac{1}{n}\left(\nu_1 + (n-1)\nu_2\right) \\
\mu_3^r =& \frac{1}{n^2}\left(\nu_1 + 3(n-1)\nu_2 + (n-1)(n-2)\nu_3\right) \\
\mu_4^r =& \frac{1}{n^3}\left(\nu_1 + 7(n-1)\nu_2 + 6(n-1)(n-2)\nu_3 + (n-1)(n-2)(n-3)\nu_4\right)
\end{aligned}
\tag{36}
$$

It is sometimes more convenient to work with central moments $\mu_j^c = E[(Z - E[Z])^j]$

$$
\begin{aligned}
\mu_1^c =& 0 \\
\mu_2^c =& \left(\frac{1}{n}\nu_1 - \nu_1^2\right) + \left(\frac{n-1}{n}\right)\nu_2 \\
\mu_3^c =& \left(\frac{1}{n^2}\nu_1 - \frac{3}{n}\nu_1^2 + 2\nu_1^3\right) + (n-1)\left(\frac{3}{n^2} - \frac{3}{n}\nu_1\right)\nu_2 + \frac{(n-1)(n-2)}{n^2}\nu_3 \\
\mu_4^c =& \left(\frac{1}{n^3}\nu_1 - \frac{4}{n^2}\nu_1^2 + \frac{6}{n}\nu_1^3 - 3\nu_1^4\right) + (n-1)\left(\frac{7}{n^3} - \frac{12}{n^2}\nu_1 + \frac{6}{n}\nu_1^2\right)\nu_2 \\
& + (n-1)(n-2)\left(\frac{6}{n^3} - \frac{4}{n^2}\nu_1\right)\nu_3 + \frac{(n-1)(n-2)(n-3)}{n^3}\nu_4
\end{aligned}
\tag{37}
$$

Furthermore the mean and CV of the underlying frequency distribution are known, so $\nu_1 = 1/S$ and $\nu_2 = (c^2+1)/S^2$ and the central moments can be written as

$$
\begin{aligned}
\mu_1^c =& 0 \\
\mu_2^c =& \left(\frac{1}{nS} - \frac{1}{S^2}\right) + \left(\frac{n-1}{n}\right)\frac{c^2+1}{S^2} \\
\mu_3^c =& \left(\frac{1}{n^2 S} - \frac{3}{nS^2} + 2\frac{1}{S^3}\right) + (n-1)\left(\frac{3}{n^2} - \frac{3}{nS}\right)\frac{c^2+1}{S^2} + \frac{(n-1)(n-2)}{n^2}\nu_3 \\
\mu_4^c =& \left(\frac{1}{n^3 S} - \frac{4}{n^2 S^2} + \frac{6}{nS^3} - \frac{3}{S^4}\right) + (n-1)\left(\frac{7}{n^3} - \frac{12}{n^2 S} + \frac{6}{nS^2}\right)\frac{c^2+1}{S^2} \\
& + (n-1)(n-2)\left(\frac{6}{n^3} - \frac{4}{n^2 S}\right)\nu_3 + \frac{(n-1)(n-2)(n-3)}{n^3}\nu_4
\end{aligned}
\tag{38}
$$

To see that the estimator $\hat{c^2}$ is unbiased, note The sample variance $V$ defined in (13) is an unbiased estimator of the variance of the compound-binomial variable, so

$$
\begin{aligned}
E[V] =& \; E\left[\left(Z_1 - \bar{Z}\right)^2\right] \\
=& \; \mu_2^c \\
=& \; \frac{S-1}{nS^2} + \frac{(n-1)}{n}\frac{c^2}{S^2}
\end{aligned}
\tag{39}
$$

The naive CV estimator $\hat{c_{\texttt{naive}}}$ introduced in section 9 is the square root of the sample variance scaled by the mean $1/S$, so it follows that

$$E[\hat{c_{\texttt{naive}}}^2] = \frac{S-1}{n} + \frac{(n-1)}{n}c^2 \tag{40}$$

In other words, the bias of $\hat{c_{\texttt{naive}}}$ is

$$\texttt{Bias}[\hat{c_{\texttt{naive}}}] = \sqrt{\frac{S-1}{n} + \frac{(n-1)}{n}c^2} - c \tag{41}$$

Rearranging, it follows that $E[\hat{c^2}] = c^2$

##### 10.6.3 Raw Moments Underlying a Compound-Binomial Distribution

If the underlying probability distribution is a beta then the raw moments $\nu_j$ are the raw moments of a beta. With the beta parameterized by its mean $1/S$ and CV $c$, the first four raw moments are

$$\begin{aligned}
\nu_1 &= \frac{1}{S} \\
\nu_2 &= \nu_1 \frac{c^2+1}{S} \\
\nu_3 &= \nu_2 \frac{(2S-1)c^2 + (S-1)}{S(S-1+c^2)} \\
\nu_4 &= \nu_3 \frac{(3S-1)c^2 + (S-1)}{S(S-1+2c^2)}
\end{aligned} \tag{42}$$

$\nu_j$ is $O(c^2/S^j)$ for $j \geq 2$.

If the parametric form of the underlying probability distribution is unknown, a Method of Moments approach would be to estimate each $\nu_j$ by $\hat{\nu_j} = \sum_i Z_i/N$.
